# mRNABench: A curated benchmark for mature mRNA property and function prediction

**DOI:** 10.1101/2025.07.05.662870

**Authors:** Ruian (Ian) Shi, Taykhoom Dalal, Philip Fradkin, Divya Koyyalagunta, Simran Chhabria, Andrew Jung, Cyrus Tam, Defne Ceyhan, Jessica Lin, Kaitlin U. Laverty, Ilyes Baali, Bo Wang, Quaid Morris

**Author notes:** Equal contribution (Ruian (Ian) Shi, Taykhoom Dalal, Philip Fradkin).

## Abstract

Messenger RNA (mRNA) is central to gene expression, and its half-life, localization, and translation efficiency drive phenotypic diversity in eukaryotic cells. While supervised learning has been used to study the mRNA regulatory code, self-supervised foundation models support a wider range of transfer learning tasks. However, the dearth of standardized benchmarks limits efforts to pinpoint the strengths of various models. Here, we present mRNABench, a benchmarking suite for mature mRNA biology, focused on human transcripts, that evaluates the representational quality of mature mRNA embeddings from self-supervised nucleotide foundation models. We curate 13 datasets and 81 prediction tasks that broadly capture salient properties of mature mRNA, and assess the performance of 29 families of nucleotide foundation models for a total of 750k experiments. Using these experiments, we study parameter scaling, correlations between sequence compressibility and performance, data-splitting strategies, and the effects of self-supervised training objective on mRNA property prediction. mRNABench can be found at: https://github.com/morrislab/mRNABench.

## 1 Introduction

Nucleotide foundation models show promise as general-purpose embedding models for RNA transcripts, offering rich representations useful for diverse RNA function prediction tasks. Despite an abundance of foundation models, and corresponding benchmarks for DNA [1, 2, 3, 4] and non-coding RNA (ncRNA) [5, 6], the modelling of messenger RNA remains underexplored.

Mature messenger RNA (mRNA) is created by splicing, which selectively retains exonic regions from a pre-mRNA, creating an mRNA splice isoform. Alternative splicing is a combinatorial process that, from a single genomic locus, can generate multiple splice isoforms depending on exon choice, each with distinct properties and functions (Figure 1). Up to 90% of human genes are alternatively spliced [7] leading to substantial diversity of gene function among cells. Splicing dysregulation is implicated in cancer [8] and other diseases [9]. More broadly, mRNA-based therapeutics such as mRNA vaccines [10] are a rapidly growing area of drug development. Capturing complex aspects of mRNA biology through representation learning could accelerate scientific and therapeutic discovery, but meaningful progress requires benchmarking that reflects the unique features and functions of mRNA.

**Figure 1:**
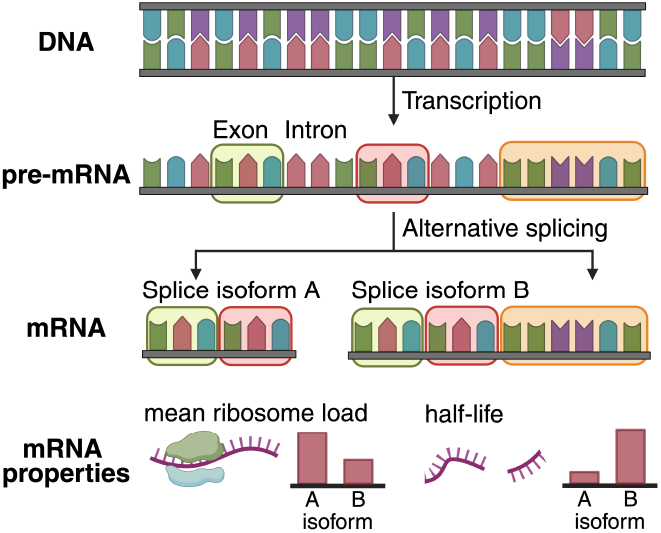
Genes are transcribed from DNA into pre-mRNA, which, through alternative splicing, can generate functionally distinct mRNA isoforms.

Current DNA and ncRNA benchmarks are not well-suited for mRNA biology. The regulatory language encoded in mRNAs is distinct from that of DNA and ncRNA, so performance on DNA and ncRNA-centric benchmarks may not translate to mRNA-based tasks. Salient mRNA functions are distinct from those evaluated in ncRNA benchmarks, and the characteristics of mRNA sequences pose unique architectural challenges. Whereas ncRNA benchmarks typically focus on short sequences in the hundreds of nucleotides and DNA benchmarks commonly operate on sub-kilobase windows of the genome, mRNA sequences regularly span kilobases. Important regulatory signal is dispersed throughout the full transcript, requiring models that integrate information across sequence scales. These factors necessitate a novel collection of tasks that assesses the representational ability of self-supervised mRNA foundation models.

To address this gap, we introduce mRNABench, a benchmarking suite with 13 distinct datasets containing 81 prediction tasks, designed to capture multiple facets of mRNA function and regulation. These tasks capture the most salient aspects of mRNA biology, including transcript stability, translational efficiency, localization, and post-transcriptional regulation. On these tasks, we use linear probing to evaluate the representational quality of 74 nucleotide foundation models, covering the majority of publicly available models, and report insights in several key areas:

### Architectural Design

We assess the impact of model size across nucleotide foundation models, and explore the suitability of contrastive learning and masked language modelling (MLM) pre-training objectives for mRNA prediction tasks whose labels arise from sequence features at different scales.

### Sequence Compressibility

We analyze the sequence content of mRNA and other genomic regions from the perspective of compression, underscoring the differences in regulatory grammar between regions. We correlate overall performance stratified by pre-training data source with these findings.

### Generalization

As genomic benchmarking often suffers from homology or sequence similarity-based data leakage [11], we use biologically-aware data-splitting strategies to assess model generalization, and report performance overestimation.

Overall, our contributions include:

- mRNABench, a Python package (https://github.com/morrislab/mRNABench) that provides lightweight access to 13 mRNA datasets with 81 function and property prediction tasks and code wrappers for 74 nucleotide foundation models. mRNABench provides integrated embedding and probing functionality, and is extensible to new datasets or models.
- Linear probe benchmarking of the above models on our curated datasets. We perform 750k experiments to assess the current state-of-the-art in mRNA modelling and analyze these results in terms of architecture choice, sequence compression, and generalization.
- A demonstration of mRNABench as a tool for model development. Through benchmarking, we show that the pre-training objective is an effective axis for model improvement alongside model scaling. Using this insight, we pre-train a joint-objective model that significantly outperforms its contrastive-only counterpart, and matches or exceeds the best existing foundation model on 6 of 13 datasets with up to ~650*×* fewer parameters.

## 2 Related Works

### Deep learning for mRNA property prediction

Supervised deep learning methods have been developed for prediction tasks such as mean ribosome loading [12, 13, 14], half-life [15, 16], subcellular localization [17, 18], expression [19], and RNA–protein interaction [20, 21], highlighting the diversity of relevant tasks in mRNA biology. Most commonly, these models use CNN-based architectures that are trained on labelled experimental data [22, 23]. While these models offer strong in-distribution prediction, they can overfit on technical noise or other dataset-specific signals. Supervised learning also typically has low sample-efficiency compared to transfer-learning approaches in mRNA property prediction [24], motivating self-supervised foundation models. We provide a detailed breakdown of all models evaluated in our benchmark in Appendix A. Several mRNA foundation models, such as mRNABERT [25], report results on mRNA tasks like mean ribosome load. However, each model is evaluated under its own protocol, preventing direct comparison between models. In mRNABench, all models are evaluated under a common probing protocol, so differences in performance reflect the learned representations instead of the evaluation setup.

### Benchmarks for Biological Sequences

Several large-scale benchmarks now exist for DNA-based tasks [1, 2, 3, 4], enabling comparisons across architectures, pre-training objectives, model sizes, etc. In contrast, few RNA-focused benchmarks have been introduced [5, 6], and they primarily evaluate RNA secondary structure and function prediction in non-coding RNAs (ncRNAs), or in zero-shot contexts [26]. Whereas ncRNA function often depends on secondary and tertiary structures [27], mRNAs are less structured [28] and are primarily regulated through linear sequence features [29, 30, 31]. As a result, benchmarks that emphasize structural accuracy fail to capture core mRNA regulatory signals. Due to the focus on ncRNA, existing benchmarks [5] focus on short sequences (hundreds of nucleotides), whereas mRNAs often span several kilobases, limiting evaluation of long-range dependencies and full-transcript representations. While long range benchmarks exist for DNA [32], the evaluated tasks do not focus on mRNA property prediction.

Existing benchmarks can also fail to account for data leakage due to homology between mRNA sequences. This can artificially inflate generalization performance, and even common strategies such as chromosomal hold-out have shown susceptibility to homology-based leakage [33]. In mRNABench, we explicitly quantify the extent of performance overestimation due to homology and evaluate how different splitting strategies affect estimates of model generalization.

## 3 Benchmarking Tasks

We curate 13 datasets, including 81 subtasks, summarized in Table 1. For each dataset, we use GenomeKit [34] to generate a six-track representation containing splice and codon positions. We further categorize our tasks as Global versus Local based on how the signal giving rise to a label is distributed in the transcript across untranslated regions (UTRs) and coding sequences (CDS). Global tasks arise primarily from aggregate effects across multiple regulatory elements, whereas Local tasks are dominated by local sequence features or perturbations (e.g., short motifs or single-nucleotide variants). While some tasks may span both regimes, this categorization provides a useful abstraction for studying how sequence information is integrated. We briefly describe each task below, and expand on data processing in Appendix B.

**Table 1:**
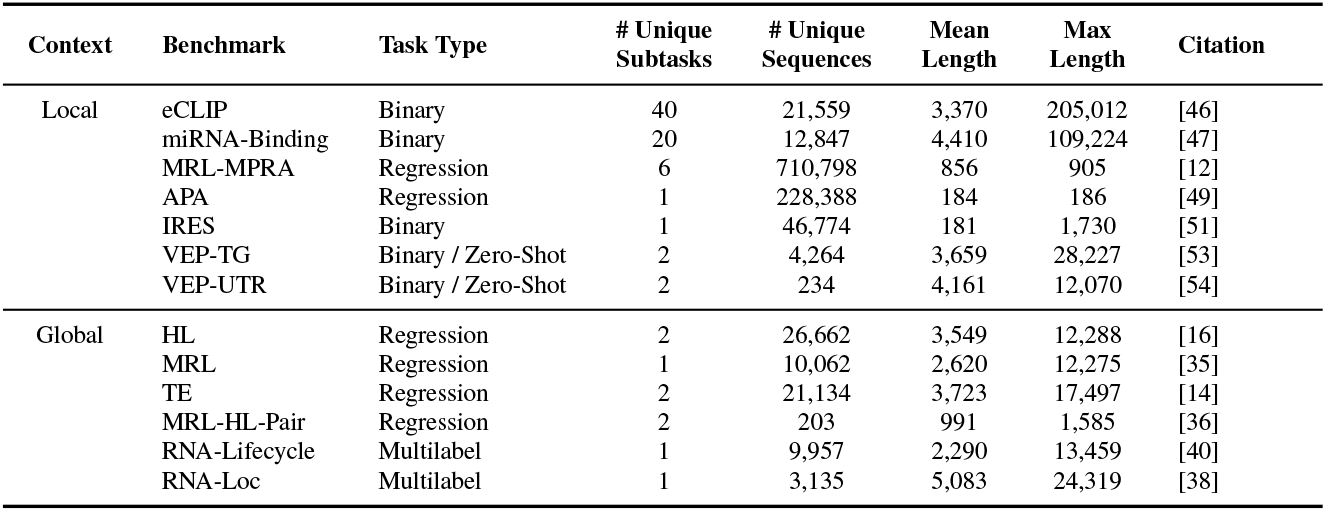
Datasets included in mRNABench, grouped by task context. For benchmarking tasks with multiple subtasks, we report the range of dataset properties. For a full description of tasks see Appendix B.

| **Context** | **Benchmark** | **Task Type** | **# Unique Subtasks** | **# Unique Sequences** | **Mean Length** | **Max Length** | **Citation** |
|---|---|---|---|---|---|---|---|
| Local | eCLIP | Binary | 40 | 21,559 | 3,370 | 205,012 | [46] |
|  | miRNA-Binding | Binary | 20 | 12,847 | 4,410 | 109,224 | [47] |
|  | MRL-MPRA | Regression | 6 | 710,798 | 856 | 905 | [12] |
|  | APA | Regression | 1 | 228,388 | 184 | 186 | [49] |
|  | IRES | Binary | 1 | 46,774 | 181 | 1,730 | [51] |
|  | VEP-TG | Binary / Zero-Shot | 2 | 4,264 | 3,659 | 28,227 | [53] |
|  | VEP-UTR | Binary / Zero-Shot | 2 | 234 | 4,161 | 12,070 | [54] |
| Global | HL | Regression | 2 | 26,662 | 3,549 | 12,288 | [16] |
|  | MRL | Regression | 1 | 10,062 | 2,620 | 12,275 | [35] |
|  | TE | Regression | 2 | 21,134 | 3,723 | 17,497 | [14] |
|  | MRL-HL-Pair | Regression | 2 | 203 | 991 | 1,585 | [36] |
|  | RNA-Lifecycle | Multilabel | 1 | 9,957 | 2,290 | 13,459 | [40] |
|  | RNA-Loc | Multilabel | 1 | 3,135 | 5,083 | 24,319 | [38] |

### 3.1 Global Tasks

**mRNA Half-Life (HL)** measures the time for half of the molecules of an mRNA transcript to degrade in the cell, and is a key determinant of transcript stability and gene expression. Longer-lived transcripts allow for more sustained protein production, whereas shorter-lived transcripts allow for rapid changes in gene expression in response to the cellular environment. We collect this data from [16], which aggregates 66 human and mouse mRNA half-life experiments.

**Mean Ribosome Load (MRL)** represents the number of ribosomes associated with an mRNA transcript, serving as a proxy for its translational efficiency. Differential translation efficiency among transcripts enables post-transcriptional regulation, as two transcripts with identical expression levels can produce vastly different amounts of protein product. Our benchmark dataset is collected from a renal cell carcinoma cell line (RCC4/VHL) [35].

**Paired mRNA Half-Life and Mean Ribosome Load (MRL-HL-Pair)** consists of synthetic mRNA sequences with paired measurements of mean ribosome load (MRL) and cellular half-life. These measurements were obtained using PERSIST-seq [36], capturing sequence variation across the 5’ UTR, CDS, and 3’ UTR, and enable joint evaluation of two key mRNA properties. With only 203 samples, this task also serves as an assessment of model performance in low-data settings.

**Translation Efficiency (TE)** uses measurements from the RiboNN dataset [14], which quantifies translational output by normalizing ribosome density to mRNA abundance. TE values are derived from thousands of ribosome profiling and RNA-seq experiments, filtered for quality based on coding region coverage. Scores are aggregated across 78 human and 68 mouse cell types, and the mean TE is used for transcript-level evaluation.

**RNA Subcellular Localization (RNA-Loc)**: The subcellular compartment to which an mRNA localizes plays a crucial role in when and where its encoded protein is synthesized [37]. We include a dataset based on APEX-seq [38], which uses an engineered enzyme to tag nearby RNAs at defined subcellular locations, enabling gene-level mapping across seven cellular compartments. We represent each gene using its APPRIS-defined principal transcript sequence [39].

**RNA Lifecycle (RNA-Lifecycle)**: This task is derived from long-read, isoform-resolved direct RNA-sequencing data from [40], which quantifies transcript flow through chromatin, cytoplasm, and polysomes. We process these data to derive multilabel transcript-level targets indicating relative depletion in each compartment, defined as transcripts falling within the bottom 33% of normalized coverage share. Depletion reflects regulatory modulation at different stages of the RNA lifecycle, including transcript processing, nuclear export, and translational initiation.

### 3.2 Local Tasks

**Massively Parallel Translation Assay - Mean Ribosome Load (MRL-MPRA)** is based on an MPRA from [12], where synthetic 5’ UTRs, either randomized or designed, were inserted upstream of reporter genes in human cells. The MRL was measured across multiple experimental conditions, including different RNA chemistries, library strategies, and reporter gene choices. Each subtask is framed as a regression predicting the measured mean ribosome load from sequence. In contrast to the previous MRL task, this task preserves the CDS while varying only the 5’ UTR. Here, translation is largely driven by local features such as the presence of upstream AUGs.

**eCLIP Binding (eCLIP)** The eCLIP protocol [41] detects the binding positions of RNA-binding proteins (RBPs). RBPs regulate mRNA processing and function co- and post-transcriptionally, through alternative splicing [42], polyadenylation [43], nuclear export [44], stability [45], among other processes. We process an eCLIP dataset using tracks collected from ENCODE [46] covering 168 RBPs across two cell lines. From this dataset, we identify the top 20 RBPs per cell line by number of bound transcripts and simplify the task for linear probing by defining a binary classification task of whether each RBP binds to a given transcript.

**miRNA Binding (miRNA-Binding)** predicts whether an mRNA transcript is targeted by specific miRNAs. We use a dataset derived from miRTarCLASH [47], which captures miRNA–mRNA interactions by sequencing chimeric miRNA–target hybrids. To mitigate class imbalance, we restrict the benchmark to the 20 most prevalent miRNAs and frame the task as a set of per-miRNA binary classification problems. An mRNA transcript is included if it is targeted by at least one miRNA.

**Alternative Polyadenylation (APA)** predicts the relative usage of competing polyadenylation sites (PAS) in the 3’ UTR, which determines 3’ UTR length and therefore the regulatory elements a transcript carries [48]. We use the APA task from BEACON [5], derived from an MPRA of synthetic 3’ UTRs containing a proximal and distal PAS [49], and predict the proportion of transcripts cleaved at the proximal site. Site choice is governed locally by the PAS motif and nearby auxiliary elements.

**IRES Classification (IRES)** predicts whether a sequence contains an internal ribosome entry site (IRES), a structured RNA element that enables translation when cap-dependent initiation is suppressed [50]. We use the binary classification dataset compiled for UTR-LM [51], which combines a bicistronic reporter screen [52] with curated IRESs from public databases. IRES activity is determined by discrete structural elements and short sequence motifs, making this a local task.

**Variant Effect Prediction - TraitGym (VEP-TG)** evaluates detection of pathogenic single nucleotide variants (SNVs) within mature mRNA transcripts. We use a filtered subset of the TraitGym dataset [53] restricted to UTR variants and retrieve the sequence of each sample using the APPRIS principal transcript [39]. Due to the restriction of sequence input to the mature mRNA region, the inherent predictability of SNV pathogenicity may be reduced.

**Variant Effect Prediction in UTR Regions (VEP-UTR)** uses a curated set of pathogenic and benign variants located in 5’ and 3’ UTRs from [54]. ClinVar variants were filtered by gnomAD allele frequency (*<* 0.05) and manually re-evaluated under ACMG guidelines to define the pathogenic set. The molecular mechanism for each variant was further examined to confirm that pathogenicity is mediated through UTR regions, and variants with confounding non-UTR effects were excluded.

## 4 Methods

### 4.1 Self-Supervised Learning Objectives

Nucleotide foundation models aim to learn useful representations of biology using self-supervised learning (SSL) on unlabelled sequence data. In this work, we focus on masked language modelling (MLM) and contrastive learning (CL), later exploring their combination during pre-training.

**Masked language modelling** is an SSL objective in which a model is trained to reconstruct masked tokens based on the surrounding sequence context. Given an input sequence *x* and a set of masked positions *M*, the MLM loss is defined as:

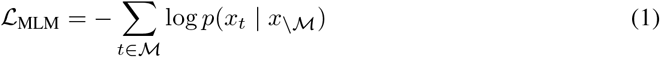

**Contrastive learning** learns global representations by bringing similar sequences (e.g., augmented views) closer in embedding space while pushing dissimilar ones apart. In this work we use the Orthrus contrastive objective [24], which applies Decoupled Contrastive Learning (DCL) [55] using splice and orthology augmentations. Given two views of the same sample 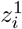 and 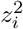 (positive pair), unrelated samples *z*_*k*_, and the temperature parameter *τ*, the DCL objective is:

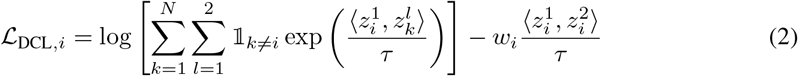

**Multi-task Objectives:** In Section 5.1, we explore combining MLM and CL using a simple multi-task objective. While the field of multi-objective optimization has proposed numerous strategies for multi-task optimization, recent work shows that simple scalarization (i.e. using fixed linear weights) is often the most effective in practice [56]. For the Orthrus+MLM model later described in Section 5.1, we use the pre-training objective: = *α*_MLM_ + (1 *α*)_CL_. Here, *α* is a weight that balances the relative contribution of each objective, set using the loss values at convergence in single-objective training so that it approximately reflects each objective’s share of the loss, and selected on validation performance (Appendix I).

### 4.2 Data Splitting Strategies

Random data splitting can overestimate model generalization due to data leakage from highly similar or homologous sequences [33]. For more rigorous assessment of generalization, mRNABench implements three biologically-informed data splitting strategies:

**Chromosomal hold-out** excludes sequences from training splits based on their chromosomal origin. While this remains a common strategy in genomics [3, 57], it assumes functional independence between chromosomes. This has been shown to cause data leakage due to sequence redundancy and homology between chromosomes [33].

**k-mer-based splitting** clusters sequences with similar short subsequence patterns (k-mers). We compute k-mer frequency vectors and cluster sequences using k-means, assigning entire clusters to a single split. This reduces leakage of low-level sequence motifs important for tasks like RNA-binding prediction, where much of the prediction performance is driven by the presence of these k-mers.

**Homology-based splitting** groups genes that share a common evolutionary origin. Using paralogous gene pairs retrieved from Ensembl Compara (release 110) [58], we apply a 35% sequence similarity threshold to filter low-confidence relationships and build transitive gene groups such that genes connected via paralogy are placed in the same data split. This is a stringent strategy to ensure that models are evaluated on unseen functional examples, though it can reduce training set diversity. We ablate the sequence similarity threshold in Appendix C.

Each method has trade-offs: chromosomal holdout is the most commonly used but most prone to data leakage, k-mer splitting reduces motif leakage but may allow some sequence-level redundancy, and homology splitting provides a stringent functional generalization test but assumes mRNA with similar functions will have similar properties. In Table 2, we use homology splitting for all datasets except for VEP, MRL-MPRA, MRL-HL-Pair, APA, and IRES, where we use random splitting. Homology splitting is unsuitable for VEP because positives and negatives are drawn from matched regions within the same gene, and for the remaining tasks because their sequences are synthetic or lack gene annotations. We evaluate the impact of each strategy in Section 5.3.

**Table 2:**
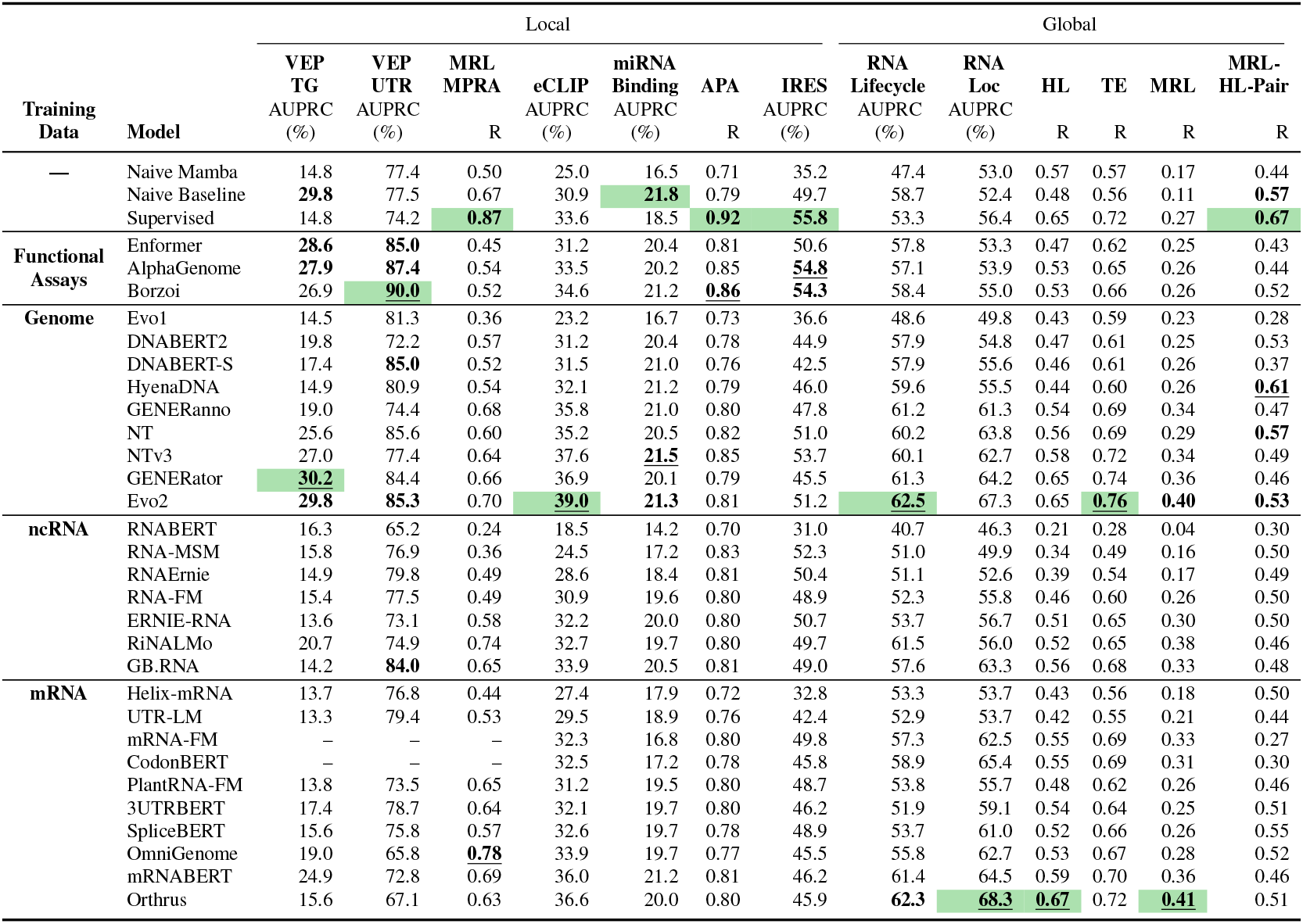
Linear probe results grouped by model category and ordered by mean of standardized metric across datasets within each category. We report the mean metric for the best model per family over ten split seeds (see Appendix G for all models). Best model for each dataset is highlighted, best foundation model is **underlined**, and models not significantly worse than it under a Wilcoxon signed-rank test (*p* = 0.05) are **bolded**.

| **Training Data** | **Model** | Local |  |  |  |  |  |  | Global |  |  |  |  |  |
|---|---|---|---|---|---|---|---|---|---|---|---|---|---|---|
|  |  | **VEP TG** | **VEP UTR** | **MRL MPRA** | **eCLIP** | **miRNA Binding** | **APA** | **IRES** | **RNA Lifecycle** | **RNA Loc** | **HL** | **TE** | **MRL** | **MRL-HL-Pair** |
|  |  | AUPRC (%) | AUPRC (%) | R | AUPRC (%) | AUPRC (%) | R | AUPRC (%) | AUPRC (%) | AUPRC (%) | R | R | R | R |
| **—** | Naive Mamba | 14.8 | 77.4 | 0.50 | 25.0 | 16.5 | 0.71 | 35.2 | 47.4 | 53.0 | 0.57 | 0.57 | 0.17 | 0.44 |
|  | Naive Baseline | **29.8** | 77.5 | 0.67 | 30.9 | **21.8** | 0.79 | 49.7 | 58.7 | 52.4 | 0.48 | 0.56 | 0.11 | **0.57** |
|  | Supervised | 14.8 | 74.2 | **0.87** | 33.6 | 18.5 | **0.92** | **55.8** | 53.3 | 56.4 | 0.65 | 0.72 | 0.27 | **0.67** |
| **Functional Assays** | Enformer | **28.6** | **85.0** | 0.45 | 31.2 | 20.4 | 0.81 | 50.6 | 57.8 | 53.3 | 0.47 | 0.62 | 0.25 | 0.43 |
|  | AlphaGenome | **27.9** | **87.4** | 0.54 | 33.5 | 20.2 | 0.85 | **54.8** | 57.1 | 53.9 | 0.53 | 0.65 | 0.26 | 0.44 |
|  | Borzoi | 26.9 | **90.0** | 0.52 | 34.6 | 21.2 | **0.86** | **54.3** | 58.4 | 55.0 | 0.53 | 0.66 | 0.26 | 0.52 |
| **Genome** | Evo1 | 14.5 | 81.3 | 0.36 | 23.2 | 16.7 | 0.73 | 36.6 | 48.6 | 49.8 | 0.43 | 0.59 | 0.23 | 0.28 |
|  | DNABERT2 | 19.8 | 72.2 | 0.57 | 31.2 | 20.4 | 0.78 | 44.9 | 57.9 | 54.8 | 0.47 | 0.61 | 0.25 | 0.53 |
|  | DNABERT-S | 17.4 | **85.0** | 0.52 | 31.5 | 21.0 | 0.76 | 42.5 | 57.9 | 55.6 | 0.46 | 0.61 | 0.26 | 0.37 |
|  | HyenaDNA | 14.9 | 80.9 | 0.54 | 32.1 | 21.2 | 0.79 | 46.0 | 59.6 | 55.5 | 0.44 | 0.60 | 0.26 | **0.61** |
|  | GENERanno | 19.0 | 74.4 | 0.68 | 35.8 | 21.0 | 0.80 | 47.8 | 61.2 | 61.3 | 0.54 | 0.69 | 0.34 | 0.47 |
|  | NT | 25.6 | 85.6 | 0.60 | 35.2 | 20.5 | 0.82 | 51.0 | 60.2 | 63.8 | 0.56 | 0.69 | 0.29 | **0.57** |
|  | NTv3 | 27.0 | 77.4 | 0.64 | 37.6 | **21.5** | 0.85 | 53.7 | 60.1 | 62.7 | 0.58 | 0.72 | 0.34 | 0.49 |
|  | GENERator | **30.2** | 84.4 | 0.66 | 36.9 | 20.1 | 0.79 | 45.5 | 61.3 | 64.2 | 0.65 | 0.74 | 0.36 | 0.46 |
|  | Evo2 | **29.8** | **85.3** | 0.70 | **39.0** | **21.3** | 0.81 | 51.2 | **62.5** | 67.3 | 0.65 | **0.76** | **0.40** | **0.53** |
| **ncRNA** | RNABERT | 16.3 | 65.2 | 0.24 | 18.5 | 14.2 | 0.70 | 31.0 | 40.7 | 46.3 | 0.21 | 0.28 | 0.04 | 0.30 |
|  | RNA-MSM | 15.8 | 76.9 | 0.36 | 24.5 | 17.2 | 0.83 | 52.3 | 51.0 | 49.9 | 0.34 | 0.49 | 0.16 | 0.50 |
|  | RNAErnie | 14.9 | 79.8 | 0.49 | 28.6 | 18.4 | 0.81 | 50.4 | 51.1 | 52.6 | 0.39 | 0.54 | 0.17 | 0.49 |
|  | RNA-FM | 15.4 | 77.5 | 0.49 | 30.9 | 19.6 | 0.80 | 48.9 | 52.3 | 55.8 | 0.46 | 0.60 | 0.26 | 0.50 |
|  | ERNIE-RNA | 13.6 | 73.1 | 0.58 | 32.2 | 20.0 | 0.80 | 50.7 | 53.7 | 56.7 | 0.51 | 0.65 | 0.30 | 0.50 |
|  | RiNALMo | 20.7 | 74.9 | 0.74 | 32.7 | 19.7 | 0.80 | 49.7 | 61.5 | 56.0 | 0.52 | 0.65 | 0.38 | 0.46 |
|  | GB.RNA | 14.2 | **84.0** | 0.65 | 33.9 | 20.5 | 0.81 | 49.0 | 57.6 | 63.3 | 0.56 | 0.68 | 0.33 | 0.48 |
| **mRNA** | Helix-mRNA | 13.7 | 76.8 | 0.44 | 27.4 | 17.9 | 0.72 | 32.8 | 53.3 | 53.7 | 0.43 | 0.56 | 0.18 | 0.50 |
|  | UTR-LM | 13.3 | 79.4 | 0.53 | 29.5 | 18.9 | 0.76 | 42.4 | 52.9 | 53.7 | 0.42 | 0.55 | 0.21 | 0.44 |
|  | mRNA-FM | – | – | – | 32.3 | 16.8 | 0.80 | 49.8 | 57.3 | 62.5 | 0.55 | 0.69 | 0.33 | 0.27 |
|  | CodonBERT | – | – | – | 32.5 | 17.2 | 0.78 | 45.8 | 58.9 | 65.4 | 0.55 | 0.69 | 0.31 | 0.30 |
|  | PlantRNA-FM | 13.8 | 73.5 | 0.65 | 31.2 | 19.5 | 0.80 | 48.7 | 53.8 | 55.7 | 0.48 | 0.62 | 0.26 | 0.46 |
|  | 3UTRBERT | 17.4 | 78.7 | 0.64 | 32.1 | 19.7 | 0.80 | 46.2 | 51.9 | 59.1 | 0.54 | 0.64 | 0.25 | 0.51 |
|  | SpliceBERT | 15.6 | 75.8 | 0.57 | 32.6 | 19.7 | 0.78 | 48.9 | 53.7 | 61.0 | 0.52 | 0.66 | 0.26 | 0.55 |
|  | OmniGenome | 19.0 | 65.8 | **0.78** | 33.9 | 19.7 | 0.77 | 45.5 | 55.8 | 62.7 | 0.53 | 0.67 | 0.28 | 0.52 |
|  | mRNABERT | 24.9 | 72.8 | 0.69 | 36.0 | 21.2 | 0.81 | 46.2 | 61.4 | 64.5 | 0.59 | 0.70 | 0.36 | 0.46 |
|  | Orthrus | 15.6 | 67.1 | 0.63 | 36.6 | 20.0 | 0.80 | 45.9 | **62.3** | **68.3** | **0.67** | 0.72 | **0.41** | 0.51 |

### 4.3 Linear Probing

Our evaluation assesses representational quality, measuring how well self-supervised pre-training produces embeddings that capture task-relevant biology. We use zero-shot and linear probing, i.e. training a linear classifier on frozen embeddings [59], to directly test whether a model produces such embeddings. While fine-tuning is used to benchmark foundation models on tasks with shorter sequences, it is unsuited for our context. Fine-tuning confounds representation quality with downstream adaptation capacity, and is infeasible for many benchmarked models, whose architectures lack the efficient attention implementations required to backpropagate through multi-kilobase sequences.

We compute transcript embeddings by averaging over per-nucleotide embeddings. For models with insufficient context windows, embeddings are computed in chunks and averaged. For VEP tasks, we compute embeddings for both wild-type and mutant sequences and use the embedding difference as features; we also report zero-shot probing results. Our experimental setup, including an ablation on embedding aggregation and chunking strategy, is detailed in Appendix D.

## 5 Results

We evaluate all foundation models, two naive baselines, and an *ab initio* supervised CNN on all tasks, and report mean linear probing performance across ten data splits in Table 2. Prediction sub-tasks were meanaggregated by source dataset, and we show the best model from each family. We report model-specific overall performance by first applying a z-score transform to all model performances within each dataset, and then taking the mean across datasets. The Fisher transform was applied to Pearson correlations prior to standardization. Full results for all models, descriptions of baselines, chosen data splits, and 95% confidence intervals are reported in Appendix A and G. We also report zero-shot performance in VEP tasks in Appendix H.

We observe that Evo2 performs the best overall, with Orthrus and Borzoi outperforming on several tasks. We explore Orthrus’ weakness in local tasks in Section 5.1. In Figure 2, we visualize the mean standardized metric as a function of model parameters, and observe a weak correlation between model size and overall performance, although stratification reveals that other factors such as pre-training data source also contribute significantly to performance. We further analyze model rankings in Appendix E, finding them robust to dropping small datasets and reweighting tasks, and per-family scaling in Appendix F, finding significant scaling trends in six of nine families.

**Figure 2:**
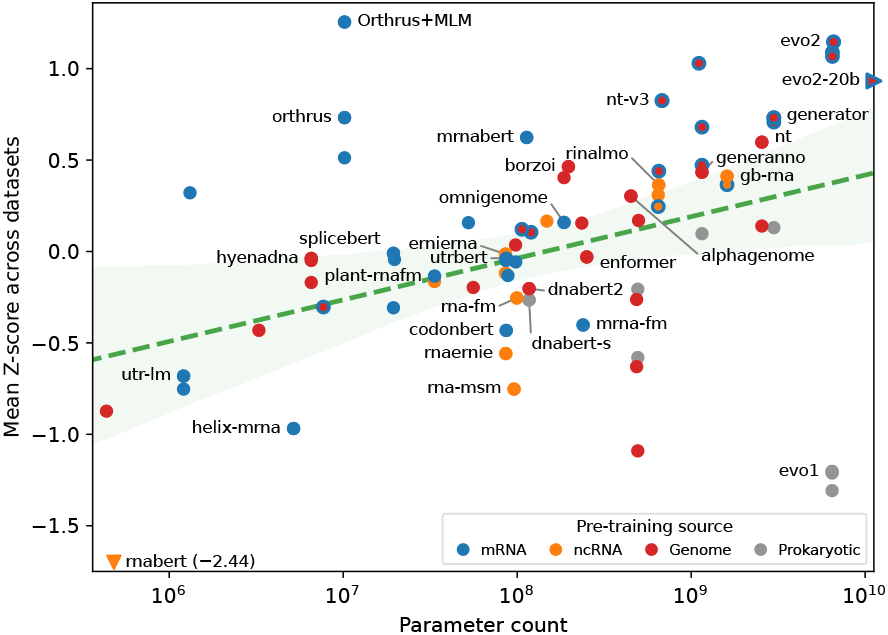
Linear probing performance. Mean of standardized metric across datasets shown on the y-axis. Data point colour shows pre-training source; ring indicates secondary source. Confidence intervals are in Appendix Table 9.

In contrast to results from the DNA foundation model space [4, 53], we find that the latest generation of foundation models outperform simpler baseline and *ab initio* supervised methods in eight of 13 tasks. While DNA language models must contend with learning from intergenic regions with low signal-to-noise ratios, the process of alternative splicing partially alleviates this issue, and consequently makes mRNA biology better suited for language-based approaches. However, we note that the regulatory language of mRNA still differs significantly from genomic DNA, and further explore this in Section 5.2.

### 5.1 Joint pre-training objectives improve mRNA foundation models

The competitive performance of the 10M parameter Orthrus model relative to the 7B parameter Evo2 model suggests that the choice of objective function plays a significant role in down-stream performance. We observe that Orthrus mainly outperforms on global tasks, consistent with findings from computer vision that contrastive pre-training yields worse performance on finer-resolution tasks [60, 61]. In Figure 3 (left), we quantify each model’s global-task bias as the difference in mean z-score between global and local tasks. Orthrus shows a strong global-task bias, consistent with this property of contrastive learning.

**Figure 3:**
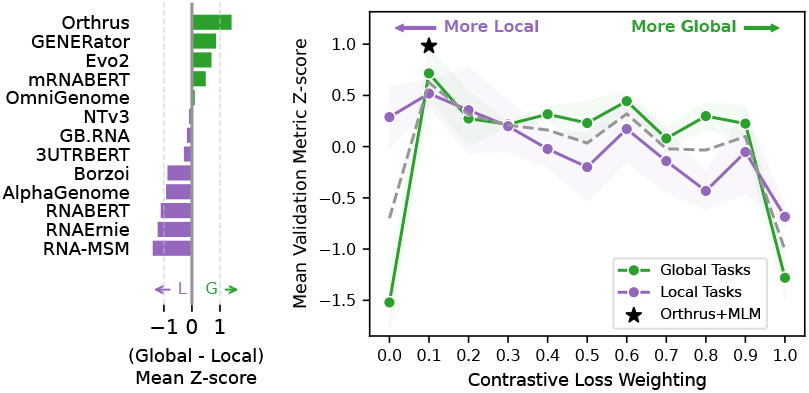
**Left:** Global-task bias (global task minus local task mean Z-score). **Right:** Joint MLM-CL model over contrastive loss weightings. Shaded area shows standard error over three seeds. Green and purple denote global and local bias (left) or tasks (right).

We hypothesize that adding an MLM loss could address this weakness in capturing local signals, and pre-train an mRNA foundation model that combines it with the Orthrus contrastive objective. We investigate the optimal ratio between these two objectives using scalarization [56]. To isolate the effect of the training objective, we fix the pre-training dataset and Mamba backbone from Orthrus and train models with varying weightings of the MLM and CL objectives. Experimental details are described in Appendix I.

Using the joint pre-training objective in Section 4.1, we trained models with CL weightings (1 *−α*) ranging from zero (MLM-only) to one (CL-only). As seen in Figure 3 (right), relying on a single objective, either MLM or CL in isolation, leads to poor overall results. Adding even a small amount of CL signal to the MLM-only model improved global task performance. At low CL weights, the model’s performance on local tasks was also unchanged, indicating that CL signal can be added to boost performance on global tasks without a corresponding drop in local task performance. Overall, we find that combining MLM and CL provides better task coverage than either alone.

We select the best-performing joint model, denoted **Orthrus+MLM**, using mean validation z-score across benchmark datasets. APA and IRES were held out from this selection as independent tasks. Orthrus+MLM beats or matches the best existing foundation model on six of 13 datasets, and significantly outperforms Orthrus on all 13, including both held-out tasks (Table 3). Contrary to our hypothesis, gains over Orthrus are similar on local and global tasks (mean z-score increase of 0.51 and 0.53, respectively), indicating that MLM improves performance overall rather than specifically addressing local tasks. Orthrus+MLM reaches the best existing model on only one of seven local tasks, compared with five of six global tasks, warranting further methodological exploration.

**Table 3:** Orthrus+MLM against its contrastive-only counterpart and the best existing foundation model (FM) per dataset, reported as in Table 2. The best row is **underlined**; rows not significantly worse than it are **bolded** (one-sided Wilcoxon signed-rank test over seeds, *p* = 0.05). Orthrus+MLM significantly outperforms Orthrus on all 13 datasets (*p <* 0.05). ^*§*^Dataset held out from Orthrus+MLM variant selection.

|  | Local |  |  |  |  |  |  | Global |  |  |  |  |  |
|---|---|---|---|---|---|---|---|---|---|---|---|---|---|
|  | **VEP<br>TG** | **VEP<br>UTR** | **MRL<br>MPRA** | **eCLIP** | **miRNA<br>Binding** | **APA**<sup>§</sup> | **IRES**<sup>§</sup> | **RNA<br>Lifecycle** | **RNA<br>Loc** | **HL** | **TE** | **MRL** | **MRL-HL-Pair** |
| Best FM | **30.2** | **90.0** | **0.78** | **39.0** | **21.5** | **0.86** | **54.8** | 62.5 | 68.3 | 0.67 | **0.76** | 0.41 | **0.61** |
| Orthrus | 15.6 | 67.1 | 0.63 | 36.6 | 20.0 | 0.80 | 45.9 | 62.3 | 68.3 | 0.67 | 0.72 | 0.41 | 0.51 |
| Orthrus+MLM | 19.5 | 83.2 | 0.64 | **39.2** | 21.1 | 0.82 | 49.7 | **64.5** | **69.4** | **0.69** | 0.75 | **0.44** | **0.63** |

### 5.2 Sequence compressibility informs model performance across genomic regions

Although nucleotide foundation models share the same input vocabulary, the sequence content and regulatory code of the genomic regions they represent differ substantially. Consequently, we expect poor performance when applying models trained on one region (e.g., the genome) to tasks in another (e.g., mRNA), a hypothesis that is supported when overall model performances are stratified by pre-training dataset source (Figure 4). Note that in Figure 4, a model is classified as “Genome only” if it never trains explicitly on full-length mRNA sequences (i.e. including UTR and CDS regions and excluding intronic regions) such as HyenaDNA [62] or Nucleotide Transformer [63], as opposed to models such as Evo2 [64] or Orthrus [24], whose pre-training data includes mRNA transcripts.

**Figure 4:**
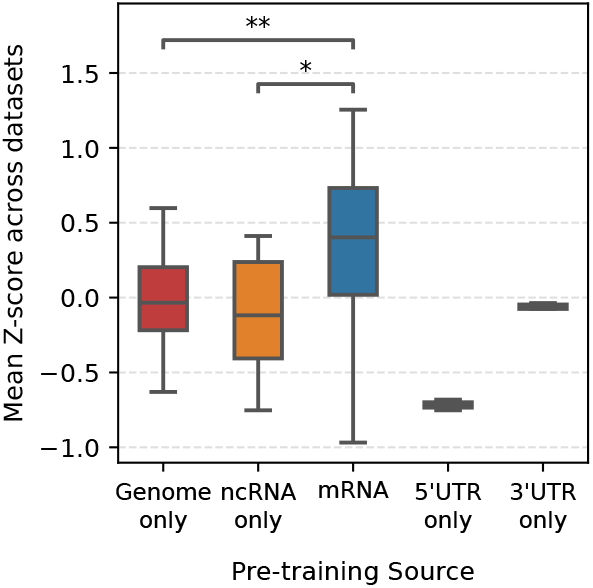
Performance stratified by pretraining data source. Significance tested using Welch’s t-test.

In this section, we aim to quantify this distributional difference among genomic regions through their compressibility. We apply a Huffman code compressor to genomic sequences, and are able to approximate the entropy of the underlying data distribution by measuring the compressed code-length [65]. In this approach, sequences with higher frequencies of occurrence are assigned shorter code lengths, thus requiring fewer bits of information for representation. Prior work has shown that DNA and RNA are far less compressible than natural language, where lower compression ratios indicate stronger statistical regularities and underlying sequence structure [66, 67].

We constructed distinct Huffman encoding schemes for various genomic regions: coding sequences (CDS), 5’ untranslated regions (UTRs), 3’ UTRs, non-coding RNA (ncRNA), intronic sequences, and intergenic DNA. 5’ UTR sequences produced the lowest compression ratio (0.951) followed by CDS (0.953), consistent with the additional structure imposed by codon usage and tightly regulated translation initiation sequences [68, 69].

We then evaluated the efficacy of each region-specific grammar in compressing data derived from the other genomic regions. A grammar obtained from CDS compressed 3’ UTR and intergenic sequences about 10% and 14% less efficiently, indicating a substantial distribution shift (Figure 5, Equation 3). In contrast, grammars originating from 3’ UTR, intergenic, intronic, and ncRNA regions compressed each other with only marginal loss in efficiency, suggesting similar sequence composition among these regions. Further analysis is described in Appendix J.

**Figure 5:**
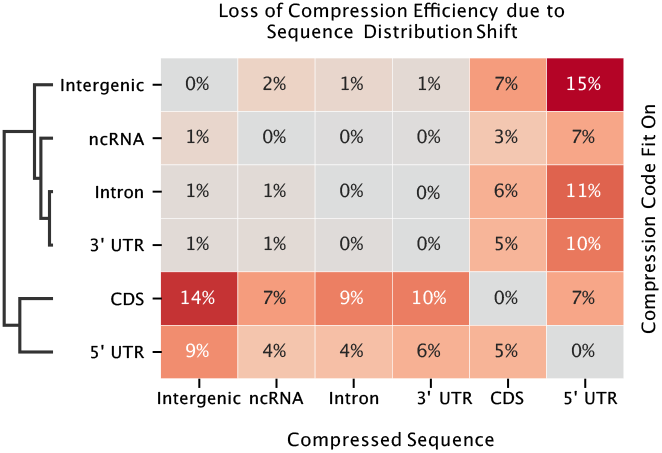
Cross-compression test scores by compression source. Numbers show percent increase in compression ratio.

These observed compression generalization gaps highlight the heterogeneity in sequence composition across different genomic regions. This analysis provides an empirical basis for understanding the generalization challenges observed when models pre-trained on non-coding sequences are applied to mRNA-specific tasks. Despite being components of mRNA, 3’ UTRs exhibit a composition resembling intergenic sequences rather than CDS and 5’ UTR regions. This distinction likely contributes to the observed model performance discrepancies.

### 5.3 Biologically-aware data splitting offers stronger estimates of generalization

One limitation in the evaluation of nucleotide foundation models lies in the overuse of naive random data splitting strategies, which tend to overestimate model generalization. This inflation arises because structurally or functionally related sequences often co-occur across data splits, inadvertently simplifying the predictive task. Given that functional outputs are frequently conserved among homologous or contextually similar sequences, models can achieve high performance by leveraging local sequence redundancy rather than by capturing broadly generalizable or mechanistically grounded features [33].

To evaluate the effect of data splitting strategy, we applied three biologically-informed splitting strategies across all applicable mRNABench tasks and compared performance to random splitting. As described in Section 4.2, we omit evaluation in VEP, MRL-MPRA, MRL-HL-Pair, APA, and IRES datasets due to their task setup, synthetic nature, or lack of gene annotations.

We visualize the results in Figure 6, where we observe that k-mer splitting consistently reduced performance. Homology-based splitting produced smaller generalization gaps, with the exception of eCLIP and MRL. Changing the homology threshold from 35% to 10% lowers performance by only 1.9% on average and leaves model rankings essentially unchanged (Spearman *ρ >* 0.99; Appendix C). We also observe trends that align with biological expectations.

**Figure 6:**
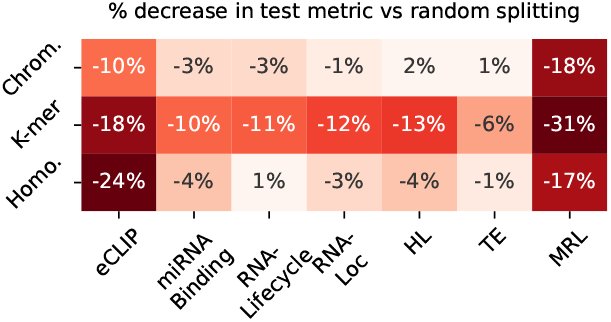
Average percent change in test metric across all models and sub-tasks compared to random splitting. All values are significant (two-sided Wilcoxon signed-rank test across models, *p <* 0.05).

For example, miRNA binding performance dropped more under k-mer splitting than homology splitting, likely because binding depends on short motifs (4–8 nucleotides) and k-mer splitting separates sequences by short-sequence composition. These results show that naive random splits overestimate generalization, and that biologically informed splits provide more realistic estimates of model robustness.

## 6 Conclusion

We present mRNABench, the first benchmarking suite focused on probing mature mRNA function and regulation. Spanning 13 distinct datasets, 81 prediction tasks, and 750k experiments, it enables standardized evaluation across 74 nucleotide foundation models, leading to three key findings:

1. Parameter scaling of nucleotide foundation models weakly correlates with mRNA task performance, though stronger trends appear within some families, and Evo2 performs best among existing models. Using benchmark-derived insights, we train a compact model with a biologically-informed joint objective that matches or exceeds state-of-the-art performance using over ~650*×* fewer parameters. This finding indicates that pre-training objective is another axis which informs model performance.
2. Models whose pre-training data includes mRNA sequences perform strongly on mRNA tasks compared to those trained on DNA or ncRNA alone, although DNA-based models remain competitive on tasks such as VEP. This is consistent with the distributional differences between genomic regions, which we quantify using compression-based analyses.
3. Naive evaluations (e.g., random splits) **overestimate generalization** and limit insight into what models have learned. Using biologically-aware splits, we show that current models fail to fully capture the modular structure of mRNA regulation.

Together, these results suggest a need to move beyond solely scaling by incorporating biological priors and designing evaluations that probe for mechanistic - rather than correlative - understanding.

### Limitations and Future Work

Several limitations of our assessment exist. The assessment omits models that performed poorly after extensive troubleshooting (Evo2 40B, reported in Appendix Table 8) [64] or had no accessible implementations [70, 71]. Many tasks are limited in the diversity of cell types, tissues, or organisms in which the property was measured, and evaluation across biological systems remains a direction for future work. One specific limitation in the VEP task lies in the choice of variants included in the dataset. While variants are filtered to be contained in UTR regions, they may act through genomic-level mechanisms (e.g. splicing or regulatory disruption), which lie outside the intended scope of this paper. We also do not control for overlap between pre-training corpora and benchmark sequences, which is likely substantial for non-synthetic tasks, although there is minimal label leakage. A detailed analysis is left for future work. In general, comparisons between models are not fully controlled, as they differ in architecture, training data, and objective. Finally, we envision future datasets on tasks such as 5’ UTR structure [28, 72] prediction will also further the completeness of our tasks. Due to the extensibility of mRNABench, we envision being able to add these datasets in future releases.

## Acknowledgments and Disclosure of Funding

We are grateful to the High Performance Computing group at Memorial Sloan Kettering Cancer Center for providing assistance, support and compute resources; this work would have been impossible without them. Resources used in preparing this research were also provided, in part, by the Province of Ontario, the Government of Canada, through the Canadian Institute for Advanced Research (CIFAR) and companies sponsoring the Vector Institute.

This work is supported by an NIH Grant to Q.M. (R01 HG013328). This research was also funded in part through the NIH/NCI Cancer Center Support Grant P30 CA008748.

R.S., P.F., and A.J. are supported by a Vector Institute Research Grant. P.F. is supported by a Natural Sciences and Engineering Research Council of Canada (NSERC) Canada Graduate Scholarship (Doctoral Program). T.D. is supported by a National Science Foundation Graduate Research Fellowship under Grant No. 2139291. D.K. is supported by a National Science Foundation Graduate Research Fellowship under Grant No. 227260-01. B.W. is supported by NSERC (grants: RGPIN-2020-06189 and DGECR-2020-00294), the Peter Munk Cardiac Centre AI Fund at the University Health Network and the CIFAR AI Chair Program.

We would like to acknowledge Stanley Z. Hua for preliminary exploration on eCLIP benchmarking.

R.S. and P.F. are co-founders of and hold equity in Blank Bio. Blank Bio did not fund this work and had no role in its design, analysis, or reporting.

## Appendix

### A Overview of Evaluated Models

All evaluated nucleotide foundation models are provided below. For each model family, written in bold, several variants are usually available, which tend to vary on model size or pre-training dataset.

| Model Name | Description | Pre-train Source | # Params (M) | Pre-train Context | Citation |
|---|---|---|---|---|---|
| **3’UTRBERT** | Transformer-based mRNA model pre-trained on 3’UTR regions of 100K human mRNA sequences. | - | - | - | [73] |
| utrbert-3mer | 3-mer tokenizer variant | mRNA | 86 | 512 |  |
| utrbert-4mer | 4-mer tokenizer variant | mRNA | 87 | 512 |  |
| utrbert-5mer | 5-mer tokenizer variant | mRNA | 88 | 512 |  |
| utrbert-6mer | 6-mer tokenizer variant | mRNA | 98 | 512 |  |
| **AlphaGenome** | Hybrid Transformer and Convolutional, U-Net style DNA sequence to function model trained to predict thousands of human and mouse genomic functional tracks at base pair resolution. | Genome | 450 | 1,000,000 | [74] |
| **Borzoi** | Hybrid Transformer and Convolutional, U-Net style DNA sequence to function model trained on human and mouse samples from ENCODE and GTEx data. The model was built to predict RNA-seq coverage from 524 kb genomic windows. | - | - | - | - |
| borzoi | - | Genome | 186 | 524,288 | [19] |
| flashzoi | Flashzoi is a variant of Borzoi trained with Flash attention. | Genome | 197 | 524,288 | [75] |
| **CodonBERT** | Transformer-based RNA model trained on 10 million mRNA CDS sequences from mammals, bacteria, and human viruses using MLM. | mRNA, Prokaryotic | 86 | 1,024 | [76] |
| **DNABERT-S** | Transformer-based DNA model trained using contrastive learning on prokaryotic genomes. | Prokaryotic | 117 | N/A | [77] |
| **DNABERT2** | Transformer-based DNA model trained on multispecies dataset. Uses BPE and other modern architectural improvements for efficiency. | Genome | 117 | N/A | [2] |
| **ERNIE-RNA** | Transformer-based RNA model trained with structural information added as attention mask biases. Pretrained on 20M ncRNA sequences. | - | - | - | [78] |
| ernie-rna | - | ncRNA | 86 | 1,024 |  |
| ernie-rna-ss | Fine-tuned on secondary structure. | ncRNA | 86 | 1,024 |  |
| **Enformer** | Hybrid Transformer and Convolutional DNA sequence to function model trained to predict over 5,000 human and 1,600 mouse functional genomic tracks such as transcriptional activity, histone modifications, TF binding, and DNA accessibility. | Genome | 251 | 196,608 | [22] |
| **Evo1** | StripedHyena-based DNA model autoregressively trained on 300b token OpenGenome dataset. | - | - | - | [79] |
| evo-1-8k-base | Base model. | Prokaryotic | 6,453 | 8,192 |  |
| evo-1-131k-base | Trained with extended context. | Prokaryotic | 6,453 | 131,072 |  |
| evo-1.5-8k-base | Extends Evo1 dataset size by 50%. | Prokaryotic | 6,453 | 8,192 |  |
| **Evo2** | StripedHyena2-based DNA model autoregressively trained on the OpenGenome2 dataset, containing 9T nucleotides across all domains of life. | - | - | - | [64] |
| evo2-1b-base | Base model trained on mixture of genomic regions including mRNA. | Genome, ncRNA, mRNA | 1,000 | 8,192 |  |
| evo2-7b-base | Base model trained on mixture of genomic regions including mRNA. | Genome, ncRNA, mRNA | 7,000 | 8,192 |  |
| evo2-7b-262k | Intermediate context model trained on long genomic sequences. | Genome, ncRNA, mRNA | 7,000 | 262,144 |  |
| evo2-7b | Extended context model trained on long genomic sequences. | Genome, ncRNA, mRNA | 7,000 | 1,000,000 |  |
| evo2-20b | Pruned from the 40B model by removing layers with high activation norms. | Genome, ncRNA, mRNA | 20,000 | 1,000,000 |  |
| evo2-40b | Despite passing author checks, we were unable to get good performance using either 40B model (40b-base and 40b). | - | - | - |  |
| **GB.RNA** | Transformer-based RNA model trained on 42M ncRNAs from RNACentral. Some variants domain adapted for mRNA using 9M coding sequences. | - | - | - | [80] |
| gb-rna-650m | - | ncRNA | 650 | 1,024 |  |
| gb-rna-1b600m | - | ncRNA | 1,600 | 1,024 |  |
| gb-rna-650m-cds | CDS adapted variant | ncRNA, mRNA | 650 | 1,024 |  |
| gb-rna-1b600m-cds | CDS adapted variant | ncRNA, mRNA | 1,600 | 1,024 |  |
| **GENERanno** | Transformer-based genomic foundation model designed for metagenomic annotation. | - | - | - | [81] |
| prokaryote-0.5b-base | Pretrained on 715 billion bp of prokaryotic DNA | Prokaryotic | 500 | 8,192 |  |
| prokaryote-0.5b-cds-annotator | Pretrained on 715 billion bp of prokaryotic DNA and finetuned for metagenomic CDS calling | Prokaryotic | 500 | 8,192 |  |
| eukaryote-0.5b-base | Pretrained on 386 billion bp of eukaryotic DNA | Genome | 500 | 8,192 |  |
| eukaryote-1.2b-cds-annotator | Pretrained on 386 billion bp of eukaryotic DNA and finetuned for metagenomic CDS calling | Genome | 1,200 | 8,192 |  |
| **GENERator** | Transformer-based autoregressive genomic foundation model trained using k-mer tokenization. The original model is trained with standard NTP on gene-centric functional regions of eukaryotic genomes. GENERator-v2 uses the same backbone and tokenization but adds Factorized Nucleotide Supervision (FNS), which decomposes each k-mer prediction into nucleotide-level likelihoods, and Genome Compression Pretraining (GCP), which concatenates functional regions. | - | - | - | - |
| eukaryote-1.2b-base | - | Genome, mRNA | 1,200 | 98,304 | [82] |
| eukaryote-3b-base | - | Genome, mRNA | 3,000 | 98,304 | [82] |
| v2-eukaryote-1.2b-base | - | Genome, mRNA | 1,200 | 98,304 | [83] |
| v2-prokaryote-1.2b-base | - | Prokaryotic | 1,200 | 98,304 | [83] |
| v2-eukaryote-3b-base | - | Genome, mRNA | 3,000 | 98,304 | [83] |
| v2-prokaryote-3b-base | - | Prokaryotic | 3,000 | 98,304 | [83] |
| **Helix-mRNA** | mRNA foundation model trained using a Mamba2 / Transformer hybrid backbone on 26M mRNAs from eukaryotic and viral species. | mRNA | 5 | 12,288 | [84] |
| **HyenaDNA** | Hyena-based DNA model trained on human reference genome using an autoregressive scheme. Variants differ in size and context length. | - | - | - | [62] |
| hyenadna-tiny-16k | - | Genome | 0.4 | 16,000 |  |
| hyenadna-small-32k | - | Genome | 3 | 32,000 |  |
| hyenadna-medium-160k | - | Genome | 7 | 160,000 |  |
| hyenadna-medium-450k | - | Genome | 7 | 450,000 |  |
| hyenadna-large-1m | - | Genome | 7 | 1,000,000 |  |
| **mRNABERT** | Transformer-based RNA foundation model trained on 36M mRNA sequences using MLM (BERT-style) pre-training. It uses ALiBi positional embeddings and Flash Attention for efficient training. It was further pre-trained using contrastive learning to align the model’s CDS embeddings with the corresponding protein embeddings from the ProtT5-XL-UniRef50 model. | mRNA | 114 | 1,024 | [25] |
| **NucleotideTransformer (NT)** | Transformer-based DNA model with variants pre-trained on differing datasets. Input tokenized to 6-mers. | - | - | - | [63] |
| 500m-human-ref | Trained on human reference. | Genome | 486 | 6,000 |  |
| 500m-1000g | Trained on 1000 Genomes Project. | Genome | 486 | 6,000 |  |
| 2.5b-1000g | Trained on 1000 Genomes Project. | Genome | 2,550 | 6,000 |  |
| 2.5b-multi-species | Trained on 850 genomes. | Genome | 2,550 | 6,000 |  |
| v2-50m-multi-species | v2 NT models use RoPE and GLUs. | Genome | 56 | 12,288 |  |
| v2-100m-multi-species | - | Genome | 98 | 12,288 |  |
| v2-250m-multi-species | - | Genome | 235 | 12,288 |  |
| v2-500m-multi-species | - | Genome | 498 | 12,288 |  |
| **NucleotideTransformerV3 (NTv3)** | Transformer/CNN-based, U-Net style DNA foundation model trained on the OpenGenome2 dataset using a masked language modelling objective at single nucleotide resolution. | - | - | - | [85] |
| v3-8M-pre | - | Genome, ncRNA, mRNA | 8 | 1,000,000 |  |
| v3-100M-pre | - | Genome, ncRNA, mRNA | 100 | 1,000,000 |  |
| v3-650M-pre | - | Genome, ncRNA, mRNA | 650 | 1,000,000 |  |
| v3-100M-post | 100M parameter model further post-trained on 15,889 functional tracks for human, mouse, fruit fly, and six plant species covering chromatin state (i.e., histone modifications) and transcriptional activity (i.e., RNA-seq) | Genome, ncRNA, mRNA | 120 | 1,000,000 |  |
| v3-650M-post | 650M parameter model further post-trained as above. | Genome, ncRNA, mRNA | 680 | 1,000,000 |  |
| **OmniGenome** | Transformer-based RNA foundation model pretrained on plant mRNA sequences from the OneKP initiative, using single-nucleotide tokenization and ViennaRNA-predicted secondary structures. | - | - | - | [86] |
| omnigenome-52m | - | mRNA | 52 | 1,024 |  |
| omnigenome-186m | - | mRNA | 186 | 1,024 |  |
| **Orthrus** | Mamba-based mRNA foundation model pre-trained on 40M mRNAs using contrastive learning on mRNA splice isoforms and orthologs. | - | - | - | [24] |
| orthrus-base-4 | - | mRNA | 1 | 12,288 |  |
| orthrus-large-4 | - | mRNA | 10 | 12,288 |  |
| orthrus-large-6 | Pre-trained using splice and codon tracks. | mRNA | 10 | 12,288 |  |
| **PlantRNA-FM** | Transformer-based RNA foundation model pretrained on 25M RNA sequences from 1,124 plant species (1KP). | mRNA | 33.5 | 1,024 | [87] |
| **RiNALMo** | Transformer-based ncRNA model trained on 36M ncRNAs. Architecture modernized to use RoPE, SwiGLU activations, and Flash Attention. | - | - | - | [88] |
| rinalmo-micro | - | ncRNA | 33 | 1,024 |  |
| rinalmo-mega | - | ncRNA | 148 | 1,024 |  |
| rinalmo-giga | - | ncRNA | 650 | 1,024 |  |
| **RNABERT** | Transformer-based ncRNA model pre-trained using both MLM and a structural alignment learning objective on 80K ncRNA sequences. | ncRNA | 0.5 | 440 | [89] |
| **RNAErnie** | Transformer-based ncRNA model pre-trained on 23M ncRNA sequences. Uses motif-level masking during pre-training. | ncRNA | 86 | 512 | [90] |
| **RNA-FM** | Transformer-based RNA foundation model pre-trained on 23M ncRNA sequences. Variant mRNA-FM trained only on CDS regions. | - | - | - | [91] |
| rna-fm | - | ncRNA | 100 | 1,024 |  |
| mrna-fm | Trained on 45M CDS-only sequences | mRNA | 239 | 1,024 |  |
| **RNA-MSM** | Transformer-based ncRNA model trained using MSA from custom structure-based homology map on roughly 8M ncRNA sequences. | ncRNA | 96 | 1,024 | [92] |
| **SpliceBERT** | Transformer-based mRNA model trained on 2M vertebrate mRNA sequences. Variants use only human RNA or use smaller contexts. | - | - | - | [93] |
| splicebert-v-1024nt | - | mRNA | 20 | 1,024 |  |
| splicebert-v-510nt | - | mRNA | 19 | 510 |  |
| splicebert-h-510nt | Trained only on human sequences. | mRNA | 19 | 510 |  |
| **UTR-LM** | Transformer-based mRNA model that is pre-trained on random and endogenous 5’UTR sequences from various species. | - | - | - | [51] |
| utrlm-te-el | Trained for translation efficiency and expression level. | mRNA | 1 | 1,026 |  |
| utrlm-mrl | Trained for mean ribosome loading. | mRNA | 1 | 1,026 |  |

Furthermore, we also implement two naive methods. NaiveBaseline uses manually extracted sequence features as embeddings. For the naive-baseline-4 model, we take all k-mers from size three to seven as features, as well as the GC content of the sequence. The naive-baseline-6 model additionally includes the CDS length and exon count as features. The NaiveMamba model is a Mamba with three layers and 64 hidden dimensions, without any pre-training (i.e., we use the ‘random’ initial weights). We note that this works surprisingly well as an embedding model in comparison to the earliest generation of nucleotide foundation models.

Finally, we include an *ab initio* supervised dilated CNN, trained from scratch on each dataset using 5 of the 10 linear probe splits (identical train, validation and test sets) and evaluated with the same metrics. The input is the one-hot sequence together with the CDS and splice tracks (6 channels). An initial convolutional layer (kernel size 3, 64 filters, BatchNorm, ReLU, MaxPool of 2) is followed by stages of dilated residual blocks. Each block has two convolutional layers (kernel size 3, stride 1, ‘same’ padding) with BatchNorm and ReLU, dropout (0.3) between them, and a residual connection; the last block of each stage applies a MaxPool of 2. During training, each sequence is shifted by a random offset of 0-3 positions. The output is mean-pooled over the sequence and passed to a two-layer head (256 hidden units).

We use a smaller model for datasets with fewer than 2,000 training examples (MRL-HL-Pair, the two UTR variant datasets and the two TraitGym datasets). It has about 270k parameters and two stages of dilated residual blocks with the following hyperparameters:

~~~
layer_params: [2, 2]
dilation_params: [1, 2]
filter_nums: [64, 128]
~~~

For all other datasets, we use a larger model of about 470k parameters with three stages:

~~~
layer_params: [2, 2, 2]
dilation_params: [1, 2, 4]
filter_nums: [64, 128, 128]
~~~

Regression targets are standardized with the training mean and standard deviation. For variant effect datasets, the model scores the variant and reference sequences and predicts their difference. eCLIP and miRNA-Binding are each trained as one multitask model per seed, and each target is reported from its own best validation step.

We train with AdamW (learning rate 0.001, weight decay 0.00001), a linear warmup over 1,000 steps followed by cosine decay, gradient clipping at 5 and bf16 mixed precision, for up to 20,000 steps. Batches group sequences of similar length, with up to 512 sequences and at most 8M padded nucleotides per batch (to avoid dropping very long transcripts). We validate every epoch (and at least once every 100 steps), stop after 1,000 steps without improvement in validation AUPRC (Pearson *r* for regression), and evaluate the best validation checkpoint. Each run used a single NVIDIA H100 GPU and took about 2-10 minutes (median 3 minutes), for a total of roughly 8 GPU-hours across all runs.

### B Dataset Processing

#### B.1 eCLIP

eCLIP data was downloaded from [46]. In total, we downloaded 244 eCLIP datasets, representing 168 unique targeted proteins; 139 of the experiments were in the K562 cell line, while the remaining 105 were in HepG2. For each eCLIP dataset, the most recent BED file was retrieved, and the corresponding BAM files from which the BED was derived were also retrieved. Given this data, we ran Peakhood [94] on both the K562 and HepG2 RBP eCLIP datasets. We retrieved a genomic annotations file (GTF) from Ensembl (release 112) [95] alongside an hg38 genomic sequences file from UCSC [96], to input as arguments to the –gtf and –gen flags, respectively. The –pre-merge flag was also used. Of the 244 total eCLIP datasets, 223 of the experiments were from a paired-end eCLIP protocol, and thus were run using only R2 reads from the eCLIP data (–bam-pp-mode 2); the remaining 21 experiments were from a single-end eCLIP protocol and were run with no BAM preprocessing (–bam-pp-mode 1). Peakhood’s output files include a set of files which identify which RNA-binding protein binding sites are associated with mature RNA transcript sequence binding (as opposed to pre-mRNA or intergenic binding sites). For this set of files, there are 2 groups, with the first group containing all possible transcript and binding site combinations and the second containing the most likely transcript sequence associated with a binding event. We chose this second group of transcript sequences as a high confidence set of sequences for the eCLIP binding task.

#### B.2 miRNA Binding

The miRNA binding dataset is derived from miRTarCLASH [47], which experimentally captures miRNA-mRNA interactions by sequencing chimeric miRNA-target hybrids in human cells. We downloaded the full human miRTarCLASH interaction table and converted it into a transcript-level binary classification benchmark. To reduce class imbalance and focus on well-supported interactions, we restricted the benchmark to the 20 most frequently observed miRNAs in the dataset. For each miRNA, we define a binary target indicating whether a given mRNA transcript is experimentally validated as being targeted by that miRNA. Transcripts not targeted by any of the selected miRNAs were excluded. Transcript sequences were constructed using GENCODE v41 annotations via GenomeKit. For each gene, we retained up to five high-priority transcripts based on annotation confidence and selected a single representative transcript per gene to avoid over-representation. Transcripts lacking gene names or not present in the reference annotation were removed. The final benchmark consists of one mature mRNA transcript per gene, each annotated with binary targets corresponding to the presence or absence of binding for each of the 20 miRNAs. Sequence inputs include the full spliced mRNA sequence along with CDS and splice-site tracks.

#### B.3 Mean Ribosome Load - MPRA

This mean ribosome load (MRL) dataset is a massively parallel reporter assay (MPRA) that assesses the translational efficiency of random and designed 5’ UTRs. The authors insert either randomized or designed 50-nucleotide sequences into a synthetic construct and measure MRL. We collected this data from the GEO repository for [12], which provides mean ribosome load measurements associated with synthetic transcripts varying in reporter gene, RNA chemistry, and 5’ UTR sequence. Specifically, the eGFP and mCherry reporters are used; three RNA chemistries are evaluated (unmodified, pseudouridine, and m1-pseudouridine); and both random and designed 5’ UTR sequences are included. We treat each unique combination of reporter gene, chemistry, and UTR design process as a separate sub-task, averaging MRL measurements across technical replicates. Sequences lacking measurements in both replicates were excluded.

#### B.4 TraitGym Variant Effect Prediction

Variant effect prediction (VEP) aims to classify single nucleotide variants (SNVs) as pathogenic or non-pathogenic. We use the TraitGym benchmark [53] and subset it to only SNVs that occur in 5’UTR and 3’UTR regions. The TraitGym benchmark matches positive and negative samples based on sequence context (e.g., distance to the transcription start site). TraitGym also stratifies SNVs into Mendelian and complex traits. We group sequences according to this TraitGym stratification and treat them as two separate sub-tasks. Unlike the original dataset, we provide embedding models with a SNV’s mature mRNA context rather than an unconstrained genomic context. To accomplish this, we searched the GENCODE v47 annotation using GenomeKit for mature mRNA transcripts that overlapped a specific SNV’s locus and used the principal transcript according to APPRIS to generate the sequence context for the SNV. This may limit the overall predictability of this task, as SNVs may cause pathogenic effects prior to splicing that are not detectable in mature mRNA. We aim to filter out these SNVs in the future.

#### B.5 5’ and 3’ UTR Variant Effect Prediction

We use the dataset from [54] which consists of 93 3’ UTR and 92 5’ UTR variants with pathogenic or benign annotations confirmed via manual literature review. To calculate variant effect, we provide the models with the variant’s mature mRNA context as in the TraitGym dataset above. We did this by retrieving the mature mRNA transcript sequence using the transcript ids provided by the authors from the NCBI RefSeq v109 annotation using GenomeKit.

#### B.6 RNA Half-life

RNA half-life measures the time elapsed for half of the molecules of a given transcript to degrade. This dataset was collected from the Saluki paper [16], which itself is an aggregate of 66 experimental datasets on human and mouse. We treat each species as a separate sub-task. Due to the noisiness of RNA half-life measurements, the authors of Saluki instead propose regressing on the first principal component of the experiment-by-gene matrix. We collect the RNA half-life dataset from the Orthrus [24] data release on Zenodo: https://zenodo.org/records/14708163. We converted the six-track encoding back into a nucleotide sequence, and use GenomeKit to map transcripts to their chromosomes using the gene name associated with each transcript. Sequences without a gene name were dropped. See the Orthrus paper for additional details on dataset composition.

#### B.7 Mean Ribosome Load

Mean ribosome load is a measure of the average number of ribosomes which are attached to a specific transcript, which is highly correlated with the translation rate of said transcript. We collect the MRL dataset from the Orthrus [24] data release on Zenodo: https://zenodo.org/records/14708163. The original data source for this dataset can be found at [35]. We converted the six-track encoding back into a nucleotide sequence, and use GenomeKit to map transcripts to their chromosomes using the gene name associated with each transcript. See the Orthrus paper for additional details on dataset composition.

#### B.8 Translation Efficiency

Translation efficiency (TE) measures protein output per mRNA molecule and reflects regulation at the level of translation. We construct human and mouse translation efficiency benchmarks using transcript-level TE measurements from [14]. For each species, we matched transcript identifiers to reference annotations using GenomeKit and excluded transcripts without valid annotations or TE measurements. For each retained transcript, we generated the full mature mRNA sequence along with coding sequence (CDS) and splice-site tracks. Translation efficiency prediction is framed as a regression task using the provided TE values as targets.

#### B.9 Paired mRNA Half-Life and MRL

We collect this data from [36], which uses PERSIST-Seq to get paired measurements of mean ribosome load and RNA half-life for a specific synthetic sequence. Also reported were various predicted measures of mRNA secondary structure, which were unused in this study but could be a future prediction target. As the source data contained sequences for the 5’UTR, CDS, and 3’UTR regions in each synthetic transcript, we were able to reconstruct the full sequence and CDS tracks.

The synthetic sequences did not undergo splicing. We drop all transcripts without both an HL and MRL measurement. During evaluation, we treat the prediction of MRL and HL as separate sub-tasks.

#### B.10 RNA Lifecycle

The dataset used in this study [40] investigates the dynamics of RNA as it is released from chromatin, exported from the nucleus, loaded onto polysomes, and degraded in both the nucleus and cytoplasm in human cells. Direct RNA sequencing data (SQK-RNA002; GEO GSE208225) comprises two replicates each of chromatin, cytoplasm, polysome and total RNA fractions from K562 cells; basecalled FASTQ files were provided by the study authors. We processed them with the `epi2me-labs/wf-transcriptomes` pipeline (v1.6.1) against the GENCODE v47 annotation and GRCh38 primary assembly; the full configuration is available in our repository. The pipeline’s differential expression output was not used. From each sample’s transcript table, we retained transcripts matching the reference intron chain, averaged StringTie coverage across replicates, and kept transcripts with positive coverage in total RNA and in at least one subcellular fraction. We normalized chromatin, cytoplasm and polysome coverage to sum to one per transcript, and defined a positive (depleted) label for a fraction when a transcript’s share is strictly below that fraction’s 33rd percentile. The final dataset has 10,043 rows covering 9,957 unique transcripts.

#### B.11 RNA Subcellular Localization

This dataset investigates the subcellular localization of mRNA transcripts in HEK293T cells using APEX-seq [38]. Following the original study’s enrichment criteria, a gene is positive for a compartment if its DESeq2 log2 fold-change over controls exceeds 0.75 with FDR-adjusted *p <* 0.05, giving a multi-label task over seven compartments: Nucleus, Nucleolus, Nuclear Lamina, Nuclear Pore, Cytosol, ER Membrane and Outer Mitochondrial Membrane. The ER lumen has no enriched transcripts and is not a label. We retain genes with estimates across all eight locations that are enriched in at least one compartment, giving 3,135 genes, each represented by its APPRIS principal transcript in GENCODE v29 (or its longest transcript if none is annotated).

#### B.12 Alternative Polyadenylation

We use the APA task from BEACON [5], derived from the synthetic 3’ UTR libraries of [49], pooling the source train, validation and test files. We remove the ten-character padding from DoubleDope sequences, leaving 228,388 unique sequences of 176–186 nt. The task is to regress the proximal isoform proportion.

#### B.13 IRES Classification

We use the dataset compiled for UTR-LM [51], consisting of 46,774 unique sequences (6–1,730 nt; median 174 nt) with 9,172 positives and 37,602 negatives. 95% of sequences come from the bicistronic reporter screen [52], with the remainder from Rfam, IRESbase, IRESite and IRESpred. We use the binary labels provided by the source (IRES_class_600).

### C Homology Threshold Ablation

Our homology split groups genes connected by paralogy above a 35% sequence similarity threshold. To test whether this threshold leaves residual leakage, we rebuilt the split at stricter thresholds of 20% and 10%, which connect more genes into shared groups, and reran linear probing for all 78 models on the 10 homology-split datasets, using mean pooling and the same ten seeds.

Stricter thresholds reduce performance only slightly (Figure 7, left). Averaged over datasets and models, the test metric falls by 1.33% at 20% and 1.93% at 10%. The decrease is consistent across models (76 and 77 of 78 models score lower at 20% and 10%, respectively; one-sided Wilcoxon signed-rank test across models, *p <* 0.001 for both), indicating that a small amount of residual leakage remains at 35%. At 10%, the largest decreases are on RNA-Lifecycle (5.1%), RNA-Loc (4.0%), HL (2.9%) and miRNA-Binding (2.6%), while performance rises on MRL (4.1%) and TE (0.6%).

**Figure 7:**
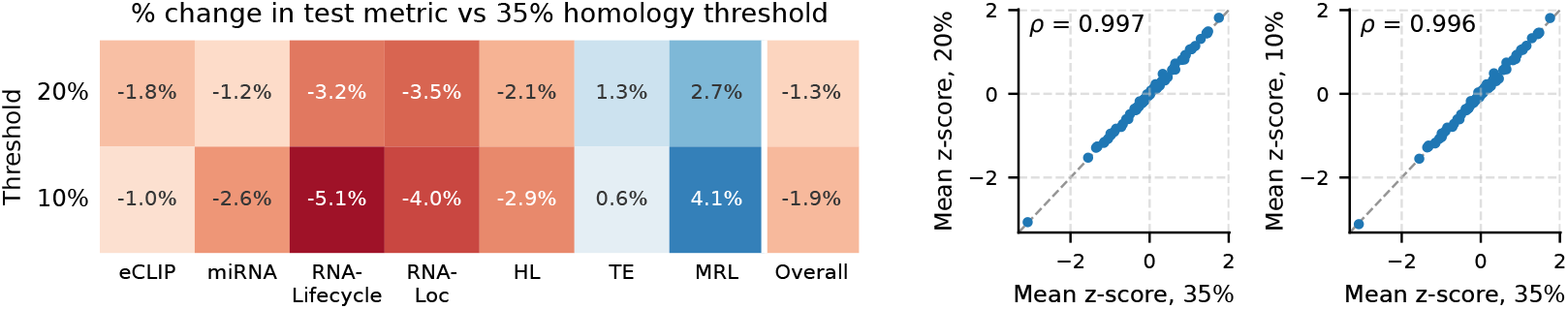
**Left:** Percent change in test metric relative to the 35% homology threshold, averaged over models for each dataset group and overall. **Right:** Mean z-score of each model at the 35% threshold against the 20% and 10% thresholds, with Spearman correlation *ρ*. The dashed line marks equal scores.

Model rankings are stable across thresholds (Figure 7, right). Spearman correlations of mean z-scores with the 35% split are 0.997 at 20% and 0.996 at 10%, and the median model moves by one rank between 35% and 10%. We therefore keep the 35% threshold, as it preserves more independent gene groups for training without changing the conclusions drawn from the benchmark.

### D Linear Probing Experimental Setup

We conduct linear probes by first extracting sequence embeddings using the nucleotide foundation models described in Appendix A. Generally, models provide a per-nucleotide or per-token embedding, and we compute the mean over the sequence length to get an *H*-dimensional vector per sequence. In the main text, we use this mean pooling for all results. However, alternative aggregations are also possible, and we assess performance when taking the max embedding across the sequence length, or concatenating the sequence mean and max embedding. These results are shown in Figure 8.

**Figure 8:**
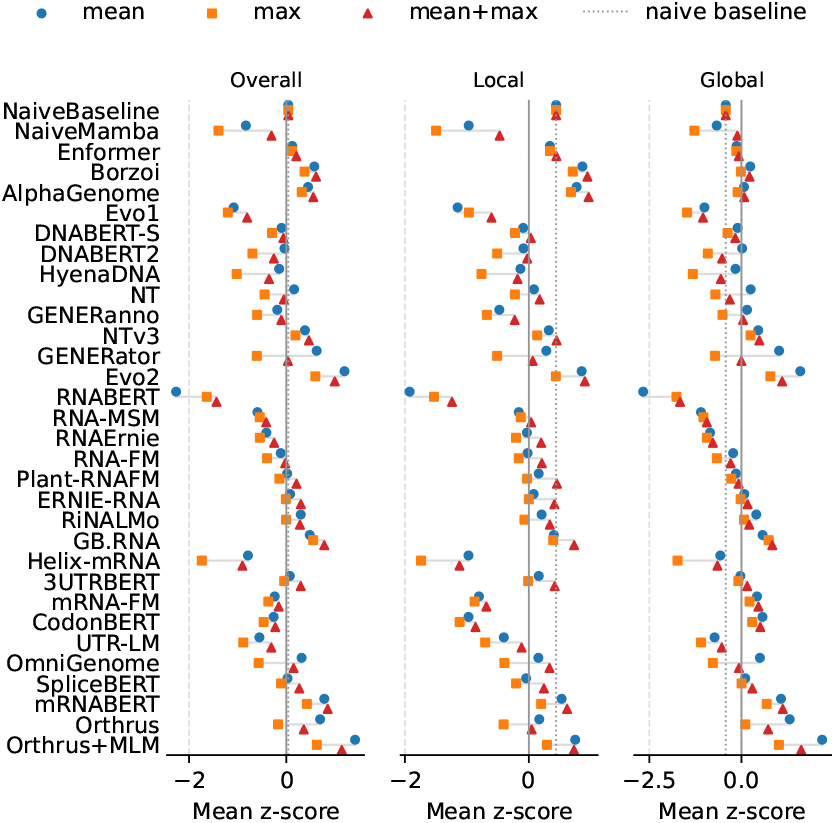
Pooling ablation. Mean z-score per model family under mean, max and mean+max pooling, overall and on local and global tasks. Z-scores are standardized jointly over the three pooling types. Naive baselines (dotted line) do not use pooling.

We find that max pooling underperforms mean pooling for almost every model, scoring higher overall for only 5 of 76 models. Mean+max is the best pooling type on local tasks for 57 of 76 models, consistent with the intuition that local tasks depend on short signals which are diluted by averaging over a long sequence. On global tasks, mean pooling is best for 47 of 76 models. Overall, rankings under mean and mean+max pooling are highly correlated (Spearman *ρ* = 0.86), and Orthrus+MLM ranks first under both. We therefore keep mean pooling as the default, while noting that mean+max is a strong choice for local tasks.

When model context length was insufficient to handle the full sequence, we computed embeddings by chunking sequences to the maximum possible length, and concatenating nucleotide embeddings across the sequence dimension prior to averaging. To test whether information is lost at boundaries when chunking, we embedded sequences with 64 or 128 nucleotides of overlap between adjacent chunks. Embeddings of overlapping positions were averaged, and the per-nucleotide embeddings were then mean-pooled as usual. We used four models with context windows of about 1,024 nucleotides (GB.RNA 650M, RiNALMo 150M, SpliceBERT-1024nt and UTR-LM) and five datasets (VEP-UTR, VEP-TG, miRNA binding, RNA-Loc and TE). In Table 4, we see that neither overlap changed performance significantly for any model.

**Table 4:** Chunk overlap ablation. Mean test metric over VEP-UTR, VEP-TG, miRNA binding, RNA-Loc and TE, averaged over 10 split seeds. *p*-values are from a paired Wilcoxon signed-rank test against no overlap, across seeds.

| Model | No overlap | 64 nt overlap (*p*) | 128 nt overlap (*p*) |
|---|---|---|---|
| GB.RNA 650M | 0.492 | 0.507 (0.105) | 0.507 (0.160) |
| RiNALMo 150M | 0.497 | 0.496 (0.275) | 0.496 (0.322) |
| SpliceBERT-1024nt | 0.503 | 0.504 (0.625) | 0.509 (0.557) |
| UTR-LM | 0.462 | 0.467 (0.322) | 0.468 (0.322) |

Once embeddings were computed for all sequences in a dataset, we perform linear probing using Scikit-Learn’s RidgeCV for regression tasks (CV performed on train split over 10 *α* values from 1e-3 to 1e6), LogisticRegressionCV for classification tasks (inverse regularization strength *C* chosen using 5-fold CV over range 1e-5 to 1e2), and a MultiOutputClassifier with LogisticRegression with above CV for multi-label tasks. The evaluation metrics are micro-averaged for multi-label tasks. For tasks (MRL) and models (NaiveBaseline) where the design matrix is too large for normal ridge regression solvers, we use the sag solver.

We replicate this linear probing across ten different data splits, using seeds [2541, 2547, 413, 412, 411, 321, 311, 421, 2515, 2516]. To aggregate performance over datasets, we first Z-transform metrics to account for differences in metric range in various tasks. Prior to standardizing Pearson correlations, we apply the Fisher transform. The Z-score is computed across models for a specific seed, dataset, and sub-task. We then mean-aggregate across sub-tasks and sub-datasets (e.g., grouping targets then cell lines for eCLIP), and then over dataset groups. We then report the mean of these values across seeds. In Appendix G, we also report the 95% confidence interval using the standard error. For results that are reported by model group, we select the model with best overall performance in the group for visual clarity.

Runtimes for linear probes were heavily dataset and model dependent. Individual linear probes can generally be run under one hour on a standard HPC CPU node, but the embedding process is highly variable. For most datasets and models, the dataset embeddings were also able to be generated within an hour on either an A100 or H100. We provide functionality within our code-base to embed sequences within a specific dataset in parallel for larger datasets.

### E Robustness of Model Rankings to Aggregation

Our overall ranking averages z-scores within dataset groups and then across groups. To test its sensitivity to this choice, we compared it in Table 5 with alternative aggregations using Spearman correlation across all models. Rankings are stable when dropping the two smallest datasets (VEP-UTR and MRL-HL-Pair) or weighting all 81 subtasks equally. Giving each category equal weight also leaves rankings largely unchanged. Rankings on individual categories diverge more, most notably on synthetic tasks (*ρ* = 0.825), which we define as MRL-MPRA, MRL-HL-Pair and APA. Table 6 shows the top ten models under each aggregation. Orthrus+MLM ranks first overall but tenth on local tasks, consistent with Section 5.1, and the supervised CNN ranks first on local tasks.

**Table 5:** Spearman correlation with the original overall ranking. Combined gives both categories in a row equal weight; First and Second only rank models using just the first or second category named in that row. Numbers in parentheses are the number of dataset groups in each category.

| **Aggregation** | **Combined** | **First only** | **Second only** |
|---|---|---|---|
| Local / Global (7 / 6) | 0.999 | 0.905 | 0.913 |
| Regression / Classification (6 / 7) | 0.999 | 0.971 | 0.925 |
| Synthetic / Natural (3 / 10) | 0.974 | 0.825 | 0.964 |
| Two smallest datasets dropped | 0.992 | – | – |
| Equal weight for all 81 subtasks | 0.967 | – | – |

**Table 6:** Top ten models under each aggregation.

| Rank | Original | Two smallest dropped | Equal subtask weight | Local only | Global only |
|---|---|---|---|---|---|
| 1 | orthrus+mlm | orthrus+mlm | evo2-7b | supervised CNN | orthrus+mlm |
| 2 | evo2-7b | evo2-7b | orthrus+mlm | nt-v3-650M-post | evo2-1b-base |
| 3 | evo2-7b-262k | evo2-7b-262k | evo2-7b-262k | evo2-7b | evo2-7b |
| 4 | evo2-7b-base | evo2-7b-base | evo2-7b-base | evo2-20b | evo2-7b-base |
| 5 | evo2-1b-base | evo2-1b-base | evo2-1b-base | evo2-7b-262k | evo2-7b-262k |
| 6 | evo2-20b | evo2-20b | nt-v3-650M-post | flashzoi | orthrus-large-6 |
| 7 | supervised CNN | nt-v3-650M-post | evo2-20b | borzoi | generator-v2-euk-3b |
| 8 | nt-v3-650M-post | supervised CNN | generator-v2-euk-3b | evo2-7b-base | orthrus-large-4 |
| 9 | orthrus-large-6 | generator-v2-euk-3b | generator-v2-euk-1.2b | alphagenome | generator-v2-euk-1.2b |
| 10 | generator-v2-euk-3b | orthrus-large-6 | mrnabert | orthrus+mlm | generator-euk-3b |

### F Parameter Scaling Within Model Families

In Figure 2, we found that the correlation between parameter count and overall performance across all models is weak. To test whether scaling helps within a model family, where other design choices are largely held fixed, we tested for a monotonic increase in performance with parameter count across the 13 dataset groups, using Page’s L trend test for families with three or more variants and a one-sided paired Wilcoxon signed-rank test otherwise. Results are shown in Table 7. Six of nine families show a significant increase (*p <* 0.05, uncorrected for multiple comparisons), indicating that scaling is effective within some families even though the correlation across all models is weak. Evo2 shows no significant trend; Evo2 40B is not included in this test.

**Table 7:** Trend in mean z-score with parameter count within model families. Δ z-score is the difference between the largest and smallest variant.

| **Family** | **Variants** | **Test** | **Δ z-score** | *p* |
|---|---|---|---|---|
| NTv2 | 4 | Page | +0.37 | <0.001 |
| NTv3 (pre) | 3 | Page | +0.74 | <0.001 |
| HyenaDNA | 3 | Page | +0.84 | <0.001 |
| RiNALMo | 3 | Page | +0.53 | <0.001 |
| GB.RNA | 2 | Wilcoxon | +0.10 | 0.013 |
| Orthrus (4-track) | 2 | Wilcoxon | +0.19 | 0.047 |
| GENERator v2 (eukaryotic) | 2 | Wilcoxon | +0.05 | 0.153 |
| OmniGenome | 2 | Wilcoxon | +0.00 | 0.446 |
| Evo2 | 3 | Page | -0.10 | 0.539 |

**Table 8:** Linear-probing results for Evo2 variants including Evo2 40B, which is left out of the main evaluation. These results predate the final dataset processing and are not comparable with Table 2; all four models share the same processing. The best variant per dataset is **bolded**. Evo2 40B scores lowest on every dataset.

| **Dataset** | **Metric** | **Evo2 1B** | **Evo2 7B** | **Evo2 20B** | **Evo2 40B** |
|---|---|---|---|---|---|
| *Local* |  |  |  |  |  |
| VEP-TG | AUPRC (%) | 26.8 | 26.1 | **30.7** | 21.0 |
| VEP-UTR | AUPRC (%) | 86.7 | 85.3 | **89.0** | 82.5 |
| MRL-MPRA | Pearson *r* | 0.666 | 0.699 | **0.720** | 0.305 |
| eCLIP | AUPRC (%) | 46.1 | **47.7** | **47.7** | 29.7 |
| miRNA-Binding | AUPRC (%) | **19.0** | 18.3 | 17.5 | 17.4 |
| *Global* |  |  |  |  |  |
| RNA-Lifecycle | AUPRC (%) | **60.7** | 59.1 | 55.6 | 49.6 |
| RNA-Loc | AUPRC (%) | **77.0** | 76.0 | 73.9 | 64.0 |
| HL | Pearson *r* | 0.633 | 0.647 | **0.651** | 0.275 |
| TE | Pearson *r* | 0.752 | **0.759** | 0.743 | 0.501 |
| MRL | Pearson *r* | 0.441 | **0.447** | 0.406 | 0.153 |
| MRL-HL-Pair | Pearson *r* | **0.572** | 0.527 | 0.528 | 0.394 |

### G Full Linear Probing Results

We report the results for each model variant, for each dataset group in Table 10 and 11. For full results, we provide a parquet file in the mRNABench GitHub repository with linear probing results for all split types, seeds, models, datasets, and tasks. Table 9 reports each model’s mean z-score, corresponding to Figure 2, overall and on local and global tasks.

**Table 9:** Mean z-score per model, ordered by overall score, with 95% confidence intervals over seeds (five seeds for the supervised CNN, ten otherwise).

| Rank | Model | Overall | Local | Global |
|---|---|---|---|---|
| 1 | orthrus+mlm | 1.25 ± 0.04 | 0.70 ± 0.08 | 1.91 ± 0.04 |
| 2 | evo2-7b | 1.15 ± 0.04 | 0.92 ± 0.06 | 1.40 ± 0.08 |
| 3 | evo2-7b-262k | 1.09 ± 0.04 | 0.85 ± 0.04 | 1.38 ± 0.09 |
| 4 | evo2-7b-base | 1.07 ± 0.03 | 0.79 ± 0.06 | 1.39 ± 0.06 |
| 5 | evo2-1b-base | 1.03 ± 0.05 | 0.65 ± 0.08 | 1.47 ± 0.07 |
| 6 | evo2-20b | 0.93 ± 0.04 | 0.86 ± 0.07 | 1.01 ± 0.06 |
| 7 | dilated-resnet | 0.92 ± 0.09 | 1.26 ± 0.15 | 0.52 ± 0.15 |
| 8 | nt-v3-650M-post | 0.83 ± 0.05 | 0.94 ± 0.09 | 0.70 ± 0.05 |
| 9 | orthrus-large-6 | 0.73 ± 0.07 | 0.18 ± 0.09 | 1.37 ± 0.07 |
| 10 | generator-v2-eukaryote-3b-base | 0.73 ± 0.05 | 0.45 ± 0.05 | 1.06 ± 0.11 |
| 11 | generator-eukaryote-3b-base | 0.71 ± 0.04 | 0.43 ± 0.05 | 1.04 ± 0.08 |
| 12 | generator-v2-eukaryote-1.2b-base | 0.68 ± 0.04 | 0.37 ± 0.08 | 1.04 ± 0.09 |
| 13 | mrnabert | 0.62 ± 0.03 | 0.47 ± 0.06 | 0.80 ± 0.05 |
| 14 | nt-2.5b-multi-species | 0.60 ± 0.04 | 0.57 ± 0.08 | 0.63 ± 0.06 |
| 15 | orthrus-large-4 | 0.51 ± 0.07 | 0.05 ± 0.13 | 1.05 ± 0.05 |
| 16 | generator-eukaryote-1.2b-base | 0.47 ± 0.03 | 0.27 ± 0.05 | 0.71 ± 0.06 |
| 17 | flashzoi | 0.46 ± 0.04 | 0.85 ± 0.05 | 0.02 ± 0.07 |
| 18 | nt-v3-650M-pre | 0.44 ± 0.04 | 0.34 ± 0.07 | 0.55 ± 0.05 |
| 19 | generanno-eukaryote-1.2b-cds-annotator-preview | 0.43 ± 0.08 | 0.34 ± 0.09 | 0.54 ± 0.08 |
| 20 | gb-rna-1b600m | 0.41 ± 0.05 | 0.52 ± 0.08 | 0.29 ± 0.07 |
| 21 | borzoi | 0.40 ± 0.07 | 0.84 ± 0.09 | -0.10 ± 0.08 |
| 22 | gb-rna-1b600m-cds | 0.37 ± 0.04 | 0.30 ± 0.06 | 0.44 ± 0.06 |
| 23 | rinalmo-giga | 0.36 ± 0.04 | 0.38 ± 0.06 | 0.34 ± 0.06 |
| 24 | orthrus-base-4 | 0.32 ± 0.06 | 0.03 ± 0.09 | 0.66 ± 0.06 |
| 25 | gb-rna-650m | 0.31 ± 0.06 | 0.39 ± 0.06 | 0.22 ± 0.06 |
| 26 | alphagenome | 0.30 ± 0.04 | 0.74 ± 0.07 | -0.21 ± 0.05 |
| 27 | gb-rna-650m-cds | 0.25 ± 0.04 | 0.15 ± 0.06 | 0.36 ± 0.07 |
| 28 | nt-v2-500m-multi-species | 0.17 ± 0.04 | 0.11 ± 0.04 | 0.24 ± 0.06 |
| 29 | rinalmo-mega | 0.17 ± 0.03 | 0.13 ± 0.05 | 0.21 ± 0.06 |
| 30 | omnigenome-186m | 0.16 ± 0.06 | 0.11 ± 0.11 | 0.22 ± 0.04 |
| 31 | omnigenome-52m | 0.16 ± 0.08 | 0.04 ± 0.11 | 0.30 ± 0.08 |
| 32 | nt-v2-250m-multi-species | 0.16 ± 0.03 | 0.12 ± 0.04 | 0.19 ± 0.03 |
| 33 | nt-2.5b-1000g | 0.14 ± 0.06 | 0.23 ± 0.09 | 0.03 ± 0.06 |
| 34 | generator-v2-prokaryote-3b-base | 0.13 ± 0.05 | -0.09 ± 0.06 | 0.39 ± 0.06 |
| 35 | nt-v3-100M-pre | 0.12 ± 0.07 | 0.11 ± 0.08 | 0.13 ± 0.09 |
| 36 | nt-v3-100M-post | 0.11 ± 0.06 | 0.30 ± 0.09 | -0.12 ± 0.08 |
| 37 | generator-v2-prokaryote-1.2b-base | 0.10 ± 0.07 | -0.07 ± 0.11 | 0.29 ± 0.06 |
| 38 | nt-v2-100m-multi-species | 0.04 ± 0.06 | 0.04 ± 0.10 | 0.03 ± 0.04 |
| 39 | splicebert-v-510nt | -0.01 ± 0.04 | -0.06 ± 0.06 | 0.05 ± 0.03 |
| 40 | ernierna | -0.01 ± 0.05 | 0.05 ± 0.07 | -0.09 ± 0.07 |
| 41 | enformer | -0.03 ± 0.05 | 0.30 ± 0.06 | -0.41 ± 0.05 |
| 42 | utrbert-4mer | -0.04 ± 0.04 | 0.07 ± 0.07 | -0.16 ± 0.05 |
| 43 | hyenadna-medium-160k | -0.04 ± 0.03 | 0.02 ± 0.05 | -0.11 ± 0.05 |
| 44 | splicebert-v-1024nt | -0.04 ± 0.05 | -0.17 ± 0.05 | 0.10 ± 0.06 |
| 45 | hyenadna-large-1m | -0.05 ± 0.05 | 0.08 ± 0.09 | -0.19 ± 0.05 |
| 46 | utrbert-3mer | -0.05 ± 0.02 | 0.06 ± 0.05 | -0.18 ± 0.02 |
| 47 | utrbert-6mer | -0.06 ± 0.03 | 0.12 ± 0.04 | -0.27 ± 0.05 |
| 48 | naive-6 | -0.06 ± 0.05 | 0.42 ± 0.08 | -0.63 ± 0.09 |
| 49 | ernierna-ss | -0.12 ± 0.04 | -0.02 ± 0.06 | -0.23 ± 0.05 |
| 50 | utrbert-5mer | -0.13 ± 0.05 | 0.10 ± 0.05 | -0.41 ± 0.05 |
| 51 | plant-rnafm | -0.14 ± 0.04 | 0.08 ± 0.06 | -0.39 ± 0.06 |
| 52 | rinalmo-micro | -0.16 ± 0.05 | -0.11 ± 0.08 | -0.23 ± 0.04 |
| 53 | naive-4 | -0.17 ± 0.04 | 0.39 ± 0.08 | -0.82 ± 0.09 |
| 54 | hyenadna-medium-450k | -0.17 ± 0.03 | -0.09 ± 0.06 | -0.27 ± 0.03 |
| 55 | nt-v2-50m-multi-species | -0.20 ± 0.04 | -0.22 ± 0.05 | -0.17 ± 0.03 |
| 56 | dnabert2 | -0.20 ± 0.05 | -0.15 ± 0.07 | -0.26 ± 0.05 |
| 57 | generanno-prokaryote-0.5b-base | -0.21 ± 0.05 | -0.48 ± 0.08 | 0.12 ± 0.07 |
| 58 | rna-fm | -0.25 ± 0.05 | -0.09 ± 0.07 | -0.45 ± 0.07 |
| 59 | nt-500m-human-ref | -0.26 ± 0.04 | -0.22 ± 0.08 | -0.31 ± 0.05 |
| 60 | dnabert-s | -0.26 ± 0.05 | -0.17 ± 0.07 | -0.38 ± 0.06 |
| 61 | nt-v3-8M-pre | -0.30 ± 0.08 | -0.36 ± 0.10 | -0.24 ± 0.09 |
| 62 | splicebert-h-510nt | -0.31 ± 0.04 | -0.11 ± 0.06 | -0.54 ± 0.05 |
| 63 | mrna-fm | -0.40 ± 0.05 | -0.88 ± 0.05 | 0.15 ± 0.08 |
| 64 | hyenadna-small-32k | -0.43 ± 0.04 | -0.37 ± 0.04 | -0.50 ± 0.06 |
| 65 | codonbert | -0.43 ± 0.04 | -1.05 ± 0.04 | 0.29 ± 0.07 |
| 66 | rnaernie | -0.56 ± 0.04 | -0.10 ± 0.07 | -1.10 ± 0.07 |
| 67 | generanno-prokaryote-0.5b-cds-annotator | -0.58 ± 0.06 | -0.88 ± 0.10 | -0.23 ± 0.08 |
| 68 | nt-500m-1000g | -0.63 ± 0.04 | -0.45 ± 0.06 | -0.84 ± 0.05 |
| 69 | utrlm-te-el | -0.68 ± 0.05 | -0.51 ± 0.08 | -0.88 ± 0.05 |
| 70 | utrlm-mrl | -0.75 ± 0.03 | -0.47 ± 0.05 | -1.08 ± 0.05 |
| 71 | rnamsm | -0.75 ± 0.05 | -0.24 ± 0.07 | -1.35 ± 0.06 |
| 72 | hyenadna-tiny-16k-d128 | -0.87 ± 0.05 | -0.69 ± 0.08 | -1.09 ± 0.05 |
| 73 | helix-mrna | -0.97 ± 0.05 | -1.09 ± 0.05 | -0.83 ± 0.11 |
| 74 | naive-mamba | -1.00 ± 0.03 | -1.10 ± 0.03 | -0.89 ± 0.06 |
| 75 | generanno-eukaryote-0.5b-base | -1.09 ± 0.04 | -1.25 ± 0.10 | -0.90 ± 0.06 |
| 76 | evo-1-131k-base | -1.20 ± 0.05 | -1.20 ± 0.08 | -1.21 ± 0.07 |
| 77 | evo-1.5-8k-base | -1.21 ± 0.03 | -1.23 ± 0.08 | -1.19 ± 0.05 |
| 78 | evo-1-8k-base | -1.31 ± 0.06 | -1.37 ± 0.09 | -1.24 ± 0.08 |
| 79 | rnabert | -2.44 ± 0.06 | -2.06 ± 0.07 | -2.88 ± 0.10 |

**Table 10:**
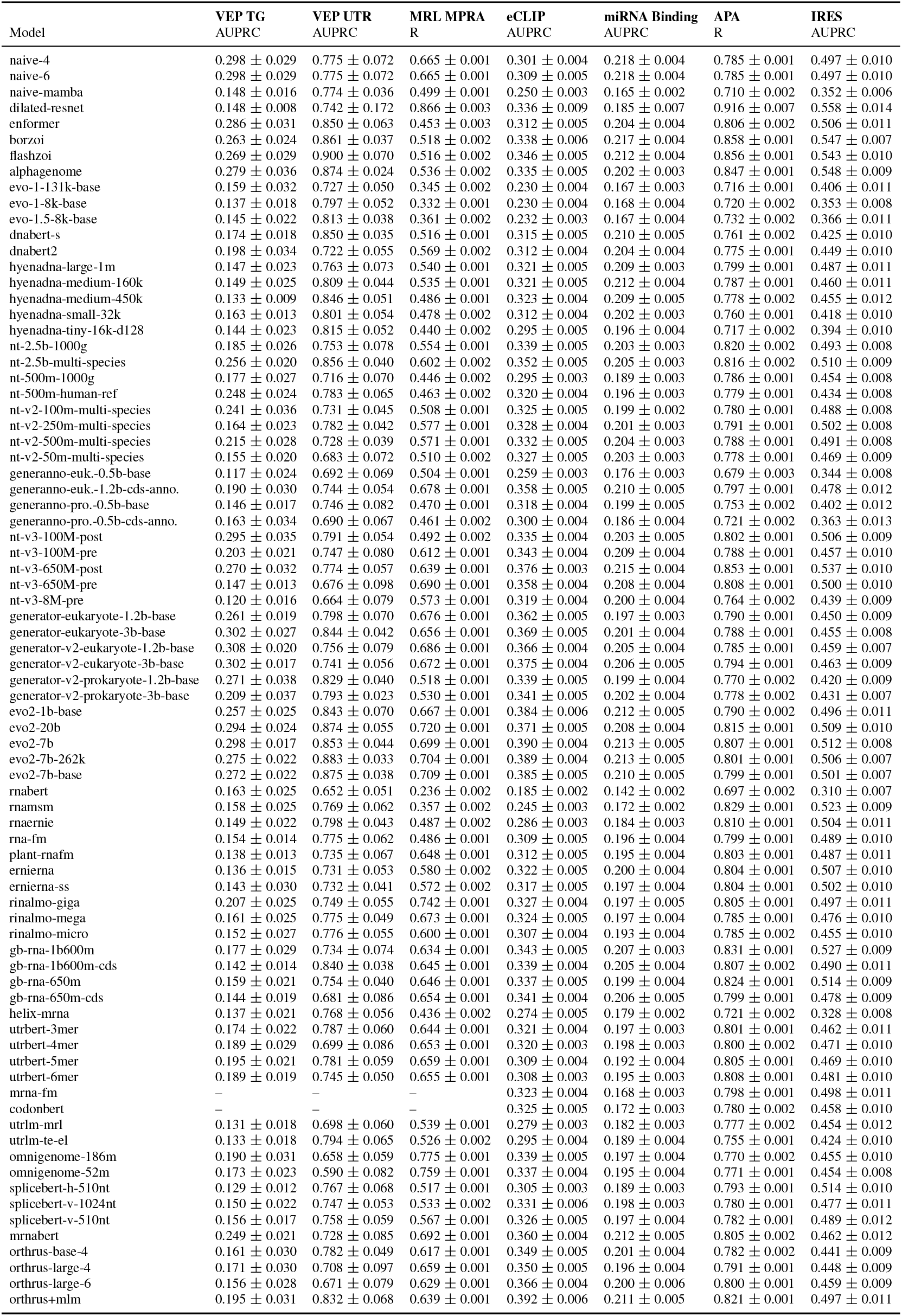
Full linear probing results. Metrics are the mean over ten data splits (five for the supervised CNN), with 95% confidence intervals. Dashes (–) indicate the embeddings were the same for every sequence.

|  | **VEP TG** | **VEP UTR** | **MRL MPRA** | **eCLIP** | **miRNA Binding** | **APA** | **IRES** |
|---|---|---|---|---|---|---|---|
| Model | AUPRC | AUPRC | R | AUPRC | AUPRC | R | AUPRC |
| naive-4 | 0.298 ± 0.029 | 0.775 ± 0.072 | 0.665 ± 0.001 | 0.301 ± 0.004 | 0.218 ± 0.004 | 0.785 ± 0.001 | 0.497 ± 0.010 |
| naive-6 | 0.298 ± 0.029 | 0.775 ± 0.072 | 0.665 ± 0.001 | 0.309 ± 0.005 | 0.218 ± 0.004 | 0.785 ± 0.001 | 0.497 ± 0.010 |
| naive-mamba | 0.148 ± 0.016 | 0.774 ± 0.036 | 0.499 ± 0.001 | 0.250 ± 0.003 | 0.165 ± 0.002 | 0.710 ± 0.002 | 0.352 ± 0.006 |
| dilated-resnet | 0.148 ± 0.008 | 0.742 ± 0.172 | 0.866 ± 0.003 | 0.336 ± 0.009 | 0.185 ± 0.007 | 0.916 ± 0.007 | 0.558 ± 0.014 |
| enformer | 0.286 ± 0.031 | 0.850 ± 0.063 | 0.453 ± 0.003 | 0.312 ± 0.005 | 0.204 ± 0.004 | 0.806 ± 0.002 | 0.506 ± 0.011 |
| borzoi | 0.263 ± 0.024 | 0.861 ± 0.037 | 0.518 ± 0.002 | 0.338 ± 0.006 | 0.217 ± 0.004 | 0.858 ± 0.001 | 0.547 ± 0.007 |
| flashzoi | 0.269 ± 0.029 | 0.900 ± 0.070 | 0.516 ± 0.002 | 0.346 ± 0.005 | 0.212 ± 0.004 | 0.856 ± 0.001 | 0.543 ± 0.010 |
| alphagenome | 0.279 ± 0.036 | 0.874 ± 0.024 | 0.536 ± 0.002 | 0.335 ± 0.005 | 0.202 ± 0.003 | 0.847 ± 0.001 | 0.548 ± 0.009 |
| evo-1-131k-base | 0.159 ± 0.032 | 0.727 ± 0.050 | 0.345 ± 0.002 | 0.230 ± 0.004 | 0.167 ± 0.003 | 0.716 ± 0.001 | 0.406 ± 0.011 |
| evo-1-8k-base | 0.137 ± 0.018 | 0.797 ± 0.052 | 0.332 ± 0.001 | 0.230 ± 0.004 | 0.168 ± 0.004 | 0.720 ± 0.002 | 0.353 ± 0.008 |
| evo-1.5-8k-base | 0.145 ± 0.022 | 0.813 ± 0.038 | 0.361 ± 0.002 | 0.232 ± 0.003 | 0.167 ± 0.004 | 0.732 ± 0.002 | 0.366 ± 0.011 |
| dnabert-s | 0.174 ± 0.018 | 0.850 ± 0.035 | 0.516 ± 0.001 | 0.315 ± 0.005 | 0.210 ± 0.005 | 0.761 ± 0.002 | 0.425 ± 0.010 |
| dnabert2 | 0.198 ± 0.034 | 0.722 ± 0.055 | 0.569 ± 0.002 | 0.312 ± 0.004 | 0.204 ± 0.004 | 0.775 ± 0.001 | 0.449 ± 0.010 |
| hyenadna-large-1m | 0.147 ± 0.023 | 0.763 ± 0.073 | 0.540 ± 0.001 | 0.321 ± 0.005 | 0.209 ± 0.003 | 0.799 ± 0.001 | 0.487 ± 0.011 |
| hyenadna-medium-160k | 0.149 ± 0.025 | 0.809 ± 0.044 | 0.535 ± 0.001 | 0.321 ± 0.005 | 0.212 ± 0.004 | 0.787 ± 0.001 | 0.460 ± 0.011 |
| hyenadna-medium-450k | 0.133 ± 0.009 | 0.846 ± 0.051 | 0.486 ± 0.001 | 0.323 ± 0.004 | 0.209 ± 0.005 | 0.778 ± 0.002 | 0.455 ± 0.012 |
| hyenadna-small-32k | 0.163 ± 0.013 | 0.801 ± 0.054 | 0.478 ± 0.002 | 0.312 ± 0.004 | 0.202 ± 0.003 | 0.760 ± 0.001 | 0.418 ± 0.010 |
| hyenadna-tiny-16k-d128 | 0.144 ± 0.023 | 0.815 ± 0.052 | 0.440 ± 0.002 | 0.295 ± 0.005 | 0.196 ± 0.004 | 0.717 ± 0.002 | 0.394 ± 0.010 |
| nt-2.5b-1000g | 0.185 ± 0.026 | 0.753 ± 0.078 | 0.554 ± 0.001 | 0.339 ± 0.005 | 0.203 ± 0.003 | 0.820 ± 0.002 | 0.493 ± 0.008 |
| nt-2.5b-multi-species | 0.256 ± 0.020 | 0.856 ± 0.040 | 0.602 ± 0.002 | 0.352 ± 0.005 | 0.205 ± 0.003 | 0.816 ± 0.002 | 0.510 ± 0.009 |
| nt-500m-1000g | 0.177 ± 0.027 | 0.716 ± 0.070 | 0.446 ± 0.002 | 0.295 ± 0.003 | 0.189 ± 0.003 | 0.786 ± 0.001 | 0.454 ± 0.008 |
| nt-500m-human-ref | 0.248 ± 0.024 | 0.783 ± 0.065 | 0.463 ± 0.002 | 0.320 ± 0.004 | 0.196 ± 0.003 | 0.779 ± 0.001 | 0.434 ± 0.008 |
| nt-v2-100m-multi-species | 0.241 ± 0.036 | 0.731 ± 0.045 | 0.508 ± 0.001 | 0.325 ± 0.005 | 0.199 ± 0.002 | 0.780 ± 0.001 | 0.488 ± 0.008 |
| nt-v2-250m-multi-species | 0.164 ± 0.023 | 0.782 ± 0.042 | 0.577 ± 0.001 | 0.328 ± 0.004 | 0.201 ± 0.003 | 0.791 ± 0.001 | 0.502 ± 0.008 |
| nt-v2-500m-multi-species | 0.215 ± 0.028 | 0.728 ± 0.039 | 0.571 ± 0.001 | 0.332 ± 0.005 | 0.204 ± 0.003 | 0.788 ± 0.001 | 0.491 ± 0.008 |
| nt-v2-50m-multi-species | 0.155 ± 0.020 | 0.683 ± 0.072 | 0.510 ± 0.002 | 0.327 ± 0.005 | 0.203 ± 0.003 | 0.778 ± 0.001 | 0.469 ± 0.009 |
| generanno-euk.-0.5b-base | 0.117 ± 0.024 | 0.692 ± 0.069 | 0.504 ± 0.001 | 0.259 ± 0.003 | 0.176 ± 0.003 | 0.679 ± 0.003 | 0.344 ± 0.008 |
| generanno-euk.-1.2b-cds-anno. | 0.190 ± 0.030 | 0.744 ± 0.054 | 0.678 ± 0.001 | 0.358 ± 0.005 | 0.210 ± 0.005 | 0.797 ± 0.001 | 0.478 ± 0.012 |
| generanno-pro.-0.5b-base | 0.146 ± 0.017 | 0.746 ± 0.082 | 0.470 ± 0.001 | 0.318 ± 0.004 | 0.199 ± 0.005 | 0.753 ± 0.002 | 0.402 ± 0.012 |
| generanno-pro.-0.5b-cds-anno. | 0.163 ± 0.034 | 0.690 ± 0.067 | 0.461 ± 0.002 | 0.300 ± 0.004 | 0.186 ± 0.004 | 0.721 ± 0.002 | 0.363 ± 0.013 |
| nt-v3-100M-post | 0.295 ± 0.035 | 0.791 ± 0.054 | 0.492 ± 0.002 | 0.335 ± 0.004 | 0.203 ± 0.005 | 0.802 ± 0.001 | 0.506 ± 0.009 |
| nt-v3-100M-pre | 0.203 ± 0.021 | 0.747 ± 0.080 | 0.612 ± 0.001 | 0.343 ± 0.004 | 0.209 ± 0.004 | 0.788 ± 0.001 | 0.457 ± 0.010 |
| nt-v3-650M-post | 0.270 ± 0.032 | 0.774 ± 0.057 | 0.639 ± 0.001 | 0.376 ± 0.003 | 0.215 ± 0.004 | 0.853 ± 0.001 | 0.537 ± 0.010 |
| nt-v3-650M-pre | 0.147 ± 0.013 | 0.676 ± 0.098 | 0.690 ± 0.001 | 0.358 ± 0.004 | 0.208 ± 0.004 | 0.808 ± 0.001 | 0.500 ± 0.010 |
| nt-v3-8M-pre | 0.120 ± 0.016 | 0.664 ± 0.079 | 0.573 ± 0.001 | 0.319 ± 0.004 | 0.200 ± 0.004 | 0.764 ± 0.002 | 0.439 ± 0.009 |
| generator-eukaryote-1.2b-base | 0.261 ± 0.019 | 0.798 ± 0.070 | 0.676 ± 0.001 | 0.362 ± 0.005 | 0.197 ± 0.003 | 0.790 ± 0.001 | 0.450 ± 0.009 |
| generator-eukaryote-3b-base | 0.302 ± 0.027 | 0.844 ± 0.042 | 0.656 ± 0.001 | 0.369 ± 0.005 | 0.201 ± 0.004 | 0.788 ± 0.001 | 0.455 ± 0.008 |
| generator-v2-eukaryote-1.2b-base | 0.308 ± 0.020 | 0.756 ± 0.079 | 0.686 ± 0.001 | 0.366 ± 0.004 | 0.205 ± 0.004 | 0.785 ± 0.001 | 0.459 ± 0.007 |
| generator-v2-eukaryote-3b-base | 0.302 ± 0.017 | 0.741 ± 0.056 | 0.672 ± 0.001 | 0.375 ± 0.004 | 0.206 ± 0.005 | 0.794 ± 0.001 | 0.463 ± 0.009 |
| generator-v2-prokaryote-1.2b-base | 0.271 ± 0.038 | 0.829 ± 0.040 | 0.518 ± 0.001 | 0.339 ± 0.005 | 0.199 ± 0.004 | 0.770 ± 0.002 | 0.420 ± 0.009 |
| generator-v2-prokaryote-3b-base | 0.209 ± 0.037 | 0.793 ± 0.023 | 0.530 ± 0.001 | 0.341 ± 0.005 | 0.202 ± 0.004 | 0.778 ± 0.002 | 0.431 ± 0.007 |
| evo2-1b-base | 0.257 ± 0.025 | 0.843 ± 0.070 | 0.667 ± 0.001 | 0.384 ± 0.006 | 0.212 ± 0.005 | 0.790 ± 0.002 | 0.496 ± 0.011 |
| evo2-20b | 0.294 ± 0.024 | 0.874 ± 0.055 | 0.720 ± 0.001 | 0.371 ± 0.005 | 0.208 ± 0.004 | 0.815 ± 0.001 | 0.509 ± 0.010 |
| evo2-7b | 0.298 ± 0.017 | 0.853 ± 0.044 | 0.699 ± 0.001 | 0.390 ± 0.004 | 0.213 ± 0.005 | 0.807 ± 0.001 | 0.512 ± 0.008 |
| evo2-7b-262k | 0.275 ± 0.022 | 0.883 ± 0.033 | 0.704 ± 0.001 | 0.389 ± 0.004 | 0.213 ± 0.005 | 0.801 ± 0.001 | 0.506 ± 0.007 |
| evo2-7b-base | 0.272 ± 0.022 | 0.875 ± 0.038 | 0.709 ± 0.001 | 0.385 ± 0.005 | 0.210 ± 0.005 | 0.799 ± 0.001 | 0.501 ± 0.007 |
| rnabert | 0.163 ± 0.025 | 0.652 ± 0.051 | 0.236 ± 0.002 | 0.185 ± 0.002 | 0.142 ± 0.002 | 0.697 ± 0.002 | 0.310 ± 0.007 |
| rnamsm | 0.158 ± 0.025 | 0.769 ± 0.062 | 0.357 ± 0.002 | 0.245 ± 0.003 | 0.172 ± 0.002 | 0.829 ± 0.001 | 0.523 ± 0.009 |
| rnaernie | 0.149 ± 0.022 | 0.798 ± 0.043 | 0.487 ± 0.002 | 0.286 ± 0.003 | 0.184 ± 0.003 | 0.810 ± 0.001 | 0.504 ± 0.011 |
| rna-fm | 0.154 ± 0.014 | 0.775 ± 0.062 | 0.486 ± 0.001 | 0.309 ± 0.005 | 0.196 ± 0.004 | 0.799 ± 0.001 | 0.489 ± 0.010 |
| plant-rnafm | 0.138 ± 0.013 | 0.735 ± 0.067 | 0.648 ± 0.001 | 0.312 ± 0.005 | 0.195 ± 0.004 | 0.803 ± 0.001 | 0.487 ± 0.011 |
| ernierna | 0.136 ± 0.015 | 0.731 ± 0.053 | 0.580 ± 0.002 | 0.322 ± 0.005 | 0.200 ± 0.004 | 0.804 ± 0.001 | 0.507 ± 0.010 |
| ernierna-ss | 0.143 ± 0.030 | 0.732 ± 0.041 | 0.572 ± 0.002 | 0.317 ± 0.005 | 0.197 ± 0.004 | 0.804 ± 0.001 | 0.502 ± 0.010 |
| rinalmo-giga | 0.207 ± 0.025 | 0.749 ± 0.055 | 0.742 ± 0.001 | 0.327 ± 0.004 | 0.197 ± 0.005 | 0.805 ± 0.001 | 0.497 ± 0.011 |
| rinalmo-mega | 0.161 ± 0.025 | 0.775 ± 0.049 | 0.673 ± 0.001 | 0.324 ± 0.005 | 0.197 ± 0.004 | 0.785 ± 0.001 | 0.476 ± 0.010 |
| rinalmo-micro | 0.152 ± 0.027 | 0.776 ± 0.055 | 0.600 ± 0.001 | 0.307 ± 0.004 | 0.193 ± 0.004 | 0.785 ± 0.002 | 0.455 ± 0.010 |
| gb-rna-1b600m | 0.177 ± 0.029 | 0.734 ± 0.074 | 0.634 ± 0.001 | 0.343 ± 0.005 | 0.207 ± 0.003 | 0.831 ± 0.001 | 0.527 ± 0.009 |
| gb-rna-1b600m-cds | 0.142 ± 0.014 | 0.840 ± 0.038 | 0.645 ± 0.001 | 0.339 ± 0.004 | 0.205 ± 0.004 | 0.807 ± 0.002 | 0.490 ± 0.011 |
| gb-rna-650m | 0.159 ± 0.021 | 0.754 ± 0.040 | 0.646 ± 0.001 | 0.337 ± 0.005 | 0.199 ± 0.004 | 0.824 ± 0.001 | 0.514 ± 0.009 |
| gb-rna-650m-cds | 0.144 ± 0.019 | 0.681 ± 0.086 | 0.654 ± 0.001 | 0.341 ± 0.004 | 0.206 ± 0.005 | 0.799 ± 0.001 | 0.478 ± 0.009 |
| helix-mrna | 0.137 ± 0.021 | 0.768 ± 0.056 | 0.436 ± 0.002 | 0.274 ± 0.005 | 0.179 ± 0.002 | 0.721 ± 0.002 | 0.328 ± 0.008 |
| utrbert-3mer | 0.174 ± 0.022 | 0.787 ± 0.060 | 0.644 ± 0.001 | 0.321 ± 0.004 | 0.197 ± 0.003 | 0.801 ± 0.001 | 0.462 ± 0.011 |
| utrbert-4mer | 0.189 ± 0.029 | 0.699 ± 0.086 | 0.653 ± 0.001 | 0.320 ± 0.003 | 0.198 ± 0.003 | 0.800 ± 0.002 | 0.471 ± 0.010 |
| utrbert-5mer | 0.195 ± 0.021 | 0.781 ± 0.059 | 0.659 ± 0.001 | 0.309 ± 0.004 | 0.192 ± 0.004 | 0.805 ± 0.001 | 0.469 ± 0.010 |
| utrbert-6mer | 0.189 ± 0.019 | 0.745 ± 0.050 | 0.655 ± 0.001 | 0.308 ± 0.003 | 0.195 ± 0.003 | 0.808 ± 0.001 | 0.481 ± 0.010 |
| mrna-fm | – | – | – | 0.323 ± 0.004 | 0.168 ± 0.003 | 0.798 ± 0.001 | 0.498 ± 0.011 |
| codonbert | – | – | – | 0.325 ± 0.005 | 0.172 ± 0.003 | 0.780 ± 0.002 | 0.458 ± 0.010 |
| utrlm-mrl | 0.131 ± 0.018 | 0.698 ± 0.060 | 0.539 ± 0.001 | 0.279 ± 0.003 | 0.182 ± 0.003 | 0.777 ± 0.002 | 0.454 ± 0.012 |
| utrlm-te-el | 0.133 ± 0.018 | 0.794 ± 0.065 | 0.526 ± 0.002 | 0.295 ± 0.004 | 0.189 ± 0.004 | 0.755 ± 0.001 | 0.424 ± 0.010 |
| omnigenome-186m | 0.190 ± 0.031 | 0.658 ± 0.059 | 0.775 ± 0.001 | 0.339 ± 0.005 | 0.197 ± 0.004 | 0.770 ± 0.002 | 0.455 ± 0.010 |
| omnigenome-52m | 0.173 ± 0.023 | 0.590 ± 0.082 | 0.759 ± 0.001 | 0.337 ± 0.004 | 0.195 ± 0.004 | 0.771 ± 0.001 | 0.454 ± 0.008 |
| splicebert-h-510nt | 0.129 ± 0.012 | 0.767 ± 0.068 | 0.517 ± 0.001 | 0.305 ± 0.003 | 0.189 ± 0.003 | 0.793 ± 0.001 | 0.514 ± 0.010 |
| splicebert-v-1024nt | 0.150 ± 0.022 | 0.747 ± 0.053 | 0.533 ± 0.002 | 0.331 ± 0.006 | 0.198 ± 0.003 | 0.780 ± 0.001 | 0.477 ± 0.011 |
| splicebert-v-510nt | 0.156 ± 0.017 | 0.758 ± 0.059 | 0.567 ± 0.001 | 0.326 ± 0.005 | 0.197 ± 0.004 | 0.782 ± 0.001 | 0.489 ± 0.012 |
| mrnabert | 0.249 ± 0.021 | 0.728 ± 0.085 | 0.692 ± 0.001 | 0.360 ± 0.004 | 0.212 ± 0.005 | 0.805 ± 0.002 | 0.462 ± 0.012 |
| orthrus-base-4 | 0.161 ± 0.030 | 0.782 ± 0.049 | 0.617 ± 0.001 | 0.349 ± 0.005 | 0.201 ± 0.004 | 0.782 ± 0.002 | 0.441 ± 0.009 |
| orthrus-large-4 | 0.171 ± 0.030 | 0.708 ± 0.097 | 0.659 ± 0.001 | 0.350 ± 0.005 | 0.196 ± 0.004 | 0.791 ± 0.001 | 0.448 ± 0.009 |
| orthrus-large-6 | 0.156 ± 0.028 | 0.671 ± 0.079 | 0.629 ± 0.001 | 0.366 ± 0.004 | 0.200 ± 0.006 | 0.800 ± 0.001 | 0.459 ± 0.009 |
| orthrus+mlm | 0.195 ± 0.031 | 0.832 ± 0.068 | 0.639 ± 0.001 | 0.392 ± 0.006 | 0.211 ± 0.005 | 0.821 ± 0.001 | 0.497 ± 0.011 |

**Table 11:**
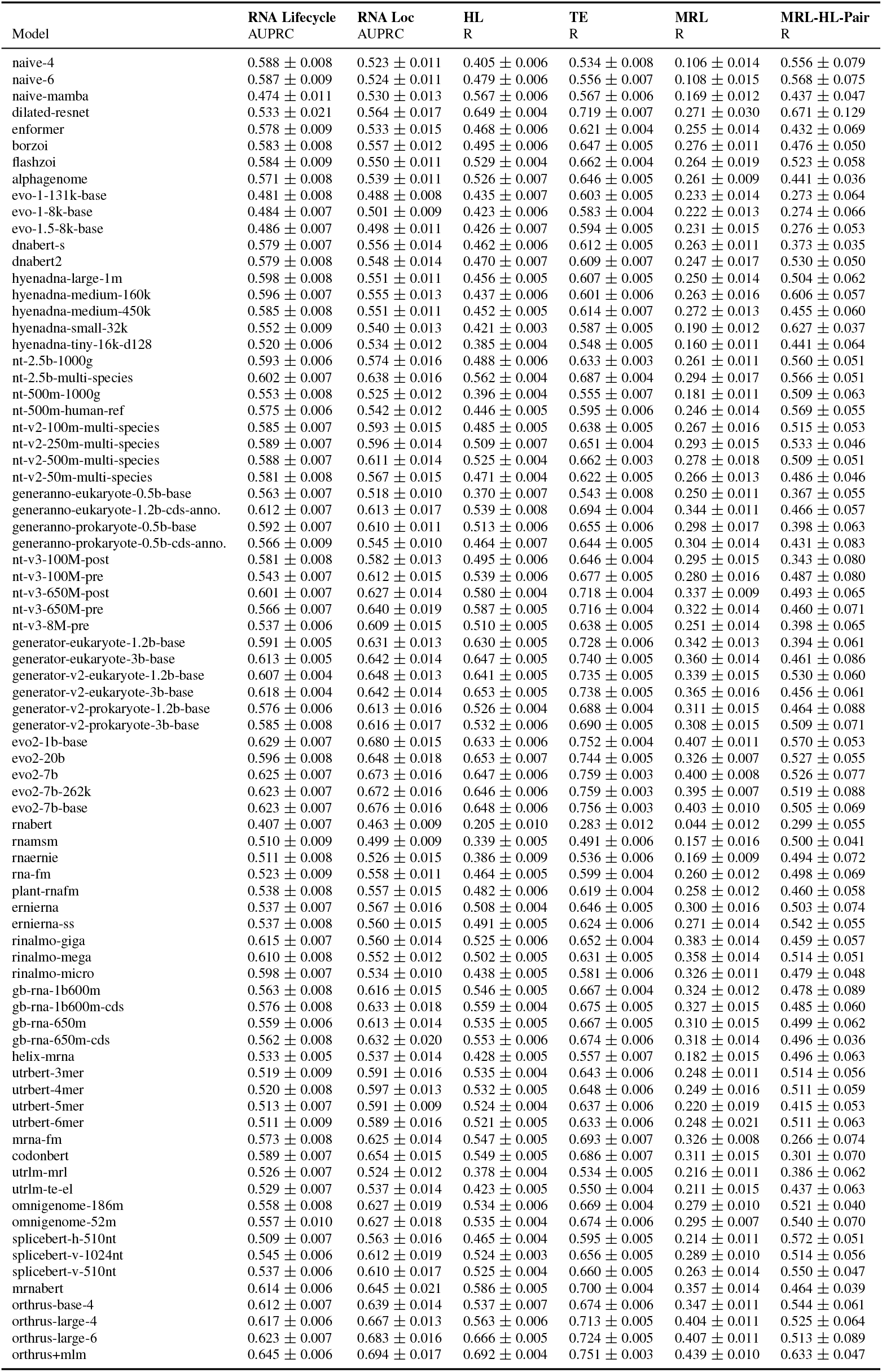
Full linear probing results. Metrics are the mean over ten data splits (five for the supervised CNN), with 95% confidence intervals.

|  | **RNA Lifecycle** | **RNA Loc** | **HL** | **TE** | **MRL** | **MRL-HL-Pair** |
|---|---|---|---|---|---|---|
| Model | AUPRC | AUPRC | R | R | R | R |
| naive-4 | 0.588 ± 0.008 | 0.523 ± 0.011 | 0.405 ± 0.006 | 0.534 ± 0.008 | 0.106 ± 0.014 | 0.556 ± 0.079 |
| naive-6 | 0.587 ± 0.009 | 0.524 ± 0.011 | 0.479 ± 0.006 | 0.556 ± 0.007 | 0.108 ± 0.015 | 0.568 ± 0.075 |
| naive-mamba | 0.474 ± 0.011 | 0.530 ± 0.013 | 0.567 ± 0.006 | 0.567 ± 0.006 | 0.169 ± 0.012 | 0.437 ± 0.047 |
| dilated-resnet | 0.533 ± 0.021 | 0.564 ± 0.017 | 0.649 ± 0.004 | 0.719 ± 0.007 | 0.271 ± 0.030 | 0.671 ± 0.129 |
| enformer | 0.578 ± 0.009 | 0.533 ± 0.015 | 0.468 ± 0.006 | 0.621 ± 0.004 | 0.255 ± 0.014 | 0.432 ± 0.069 |
| borzoi | 0.583 ± 0.008 | 0.557 ± 0.012 | 0.495 ± 0.006 | 0.647 ± 0.005 | 0.276 ± 0.011 | 0.476 ± 0.050 |
| flashzoi | 0.584 ± 0.009 | 0.550 ± 0.011 | 0.529 ± 0.004 | 0.662 ± 0.004 | 0.264 ± 0.019 | 0.523 ± 0.058 |
| alphagenome | 0.571 ± 0.008 | 0.539 ± 0.011 | 0.526 ± 0.007 | 0.646 ± 0.005 | 0.261 ± 0.009 | 0.441 ± 0.036 |
| evo-1-131k-base | 0.481 ± 0.008 | 0.488 ± 0.008 | 0.435 ± 0.007 | 0.603 ± 0.005 | 0.233 ± 0.014 | 0.273 ± 0.064 |
| evo-1-8k-base | 0.484 ± 0.007 | 0.501 ± 0.009 | 0.423 ± 0.006 | 0.583 ± 0.004 | 0.222 ± 0.013 | 0.274 ± 0.066 |
| evo-1.5-8k-base | 0.486 ± 0.007 | 0.498 ± 0.011 | 0.426 ± 0.007 | 0.594 ± 0.005 | 0.231 ± 0.015 | 0.276 ± 0.053 |
| dnabert-s | 0.579 ± 0.007 | 0.556 ± 0.014 | 0.462 ± 0.006 | 0.612 ± 0.005 | 0.263 ± 0.011 | 0.373 ± 0.035 |
| dnabert2 | 0.579 ± 0.008 | 0.548 ± 0.014 | 0.470 ± 0.007 | 0.609 ± 0.007 | 0.247 ± 0.017 | 0.530 ± 0.050 |
| hyenadna-large-1m | 0.598 ± 0.008 | 0.551 ± 0.011 | 0.456 ± 0.005 | 0.607 ± 0.005 | 0.250 ± 0.014 | 0.504 ± 0.062 |
| hyenadna-medium-160k | 0.596 ± 0.007 | 0.555 ± 0.013 | 0.437 ± 0.006 | 0.601 ± 0.006 | 0.263 ± 0.016 | 0.606 ± 0.057 |
| hyenadna-medium-450k | 0.585 ± 0.008 | 0.551 ± 0.011 | 0.452 ± 0.005 | 0.614 ± 0.007 | 0.272 ± 0.013 | 0.455 ± 0.060 |
| hyenadna-small-32k | 0.552 ± 0.009 | 0.540 ± 0.013 | 0.421 ± 0.003 | 0.587 ± 0.005 | 0.190 ± 0.012 | 0.627 ± 0.037 |
| hyenadna-tiny-16k-d128 | 0.520 ± 0.006 | 0.534 ± 0.012 | 0.385 ± 0.004 | 0.548 ± 0.005 | 0.160 ± 0.011 | 0.441 ± 0.064 |
| nt-2.5b-1000g | 0.593 ± 0.006 | 0.574 ± 0.016 | 0.488 ± 0.006 | 0.633 ± 0.003 | 0.261 ± 0.011 | 0.560 ± 0.051 |
| nt-2.5b-multi-species | 0.602 ± 0.007 | 0.638 ± 0.016 | 0.562 ± 0.004 | 0.687 ± 0.004 | 0.294 ± 0.017 | 0.566 ± 0.051 |
| nt-500m-1000g | 0.553 ± 0.008 | 0.525 ± 0.012 | 0.396 ± 0.004 | 0.555 ± 0.007 | 0.181 ± 0.011 | 0.509 ± 0.063 |
| nt-500m-human-ref | 0.575 ± 0.006 | 0.542 ± 0.012 | 0.446 ± 0.005 | 0.595 ± 0.006 | 0.246 ± 0.014 | 0.569 ± 0.055 |
| nt-v2-100m-multi-species | 0.585 ± 0.007 | 0.593 ± 0.015 | 0.485 ± 0.005 | 0.638 ± 0.005 | 0.267 ± 0.016 | 0.515 ± 0.053 |
| nt-v2-250m-multi-species | 0.589 ± 0.007 | 0.596 ± 0.014 | 0.509 ± 0.007 | 0.651 ± 0.004 | 0.293 ± 0.015 | 0.533 ± 0.046 |
| nt-v2-500m-multi-species | 0.588 ± 0.007 | 0.611 ± 0.014 | 0.525 ± 0.004 | 0.662 ± 0.003 | 0.278 ± 0.018 | 0.509 ± 0.051 |
| nt-v2-50m-multi-species | 0.581 ± 0.008 | 0.567 ± 0.015 | 0.471 ± 0.004 | 0.622 ± 0.005 | 0.266 ± 0.013 | 0.486 ± 0.046 |
| generanno-eukaryote-0.5b-base | 0.563 ± 0.007 | 0.518 ± 0.010 | 0.370 ± 0.007 | 0.543 ± 0.008 | 0.250 ± 0.011 | 0.367 ± 0.055 |
| generanno-eukaryote-1.2b-cds-anno. | 0.612 ± 0.007 | 0.613 ± 0.017 | 0.539 ± 0.008 | 0.694 ± 0.004 | 0.344 ± 0.011 | 0.466 ± 0.057 |
| generanno-prokaryote-0.5b-base | 0.592 ± 0.007 | 0.610 ± 0.011 | 0.513 ± 0.006 | 0.655 ± 0.006 | 0.298 ± 0.017 | 0.398 ± 0.063 |
| generanno-prokaryote-0.5b-cds-anno. | 0.566 ± 0.009 | 0.545 ± 0.010 | 0.464 ± 0.007 | 0.644 ± 0.005 | 0.304 ± 0.014 | 0.431 ± 0.083 |
| nt-v3-100M-post | 0.581 ± 0.008 | 0.582 ± 0.013 | 0.495 ± 0.006 | 0.646 ± 0.004 | 0.295 ± 0.015 | 0.343 ± 0.080 |
| nt-v3-100M-pre | 0.543 ± 0.007 | 0.612 ± 0.015 | 0.539 ± 0.006 | 0.677 ± 0.005 | 0.280 ± 0.016 | 0.487 ± 0.080 |
| nt-v3-650M-post | 0.601 ± 0.007 | 0.627 ± 0.014 | 0.580 ± 0.004 | 0.718 ± 0.004 | 0.337 ± 0.009 | 0.493 ± 0.065 |
| nt-v3-650M-pre | 0.566 ± 0.007 | 0.640 ± 0.019 | 0.587 ± 0.005 | 0.716 ± 0.004 | 0.322 ± 0.014 | 0.460 ± 0.071 |
| nt-v3-8M-pre | 0.537 ± 0.006 | 0.609 ± 0.015 | 0.510 ± 0.005 | 0.638 ± 0.005 | 0.251 ± 0.014 | 0.398 ± 0.065 |
| generator-eukaryote-1.2b-base | 0.591 ± 0.005 | 0.631 ± 0.013 | 0.630 ± 0.005 | 0.728 ± 0.006 | 0.342 ± 0.013 | 0.394 ± 0.061 |
| generator-eukaryote-3b-base | 0.613 ± 0.005 | 0.642 ± 0.014 | 0.647 ± 0.005 | 0.740 ± 0.005 | 0.360 ± 0.014 | 0.461 ± 0.086 |
| generator-v2-eukaryote-1.2b-base | 0.607 ± 0.004 | 0.648 ± 0.013 | 0.641 ± 0.005 | 0.735 ± 0.005 | 0.339 ± 0.015 | 0.530 ± 0.060 |
| generator-v2-eukaryote-3b-base | 0.618 ± 0.004 | 0.642 ± 0.014 | 0.653 ± 0.005 | 0.738 ± 0.005 | 0.365 ± 0.016 | 0.456 ± 0.061 |
| generator-v2-prokaryote-1.2b-base | 0.576 ± 0.006 | 0.613 ± 0.016 | 0.526 ± 0.004 | 0.688 ± 0.004 | 0.311 ± 0.015 | 0.464 ± 0.088 |
| generator-v2-prokaryote-3b-base | 0.585 ± 0.008 | 0.616 ± 0.017 | 0.532 ± 0.006 | 0.690 ± 0.005 | 0.308 ± 0.015 | 0.509 ± 0.071 |
| evo2-1b-base | 0.629 ± 0.007 | 0.680 ± 0.015 | 0.633 ± 0.006 | 0.752 ± 0.004 | 0.407 ± 0.011 | 0.570 ± 0.053 |
| evo2-20b | 0.596 ± 0.008 | 0.648 ± 0.018 | 0.653 ± 0.007 | 0.744 ± 0.005 | 0.326 ± 0.007 | 0.527 ± 0.055 |
| evo2-7b | 0.625 ± 0.007 | 0.673 ± 0.016 | 0.647 ± 0.006 | 0.759 ± 0.003 | 0.400 ± 0.008 | 0.526 ± 0.077 |
| evo2-7b-262k | 0.623 ± 0.007 | 0.672 ± 0.016 | 0.646 ± 0.006 | 0.759 ± 0.003 | 0.395 ± 0.007 | 0.519 ± 0.088 |
| evo2-7b-base | 0.623 ± 0.007 | 0.676 ± 0.016 | 0.648 ± 0.006 | 0.756 ± 0.003 | 0.403 ± 0.010 | 0.505 ± 0.069 |
| rnabert | 0.407 ± 0.007 | 0.463 ± 0.009 | 0.205 ± 0.010 | 0.283 ± 0.012 | 0.044 ± 0.012 | 0.299 ± 0.055 |
| rnamsm | 0.510 ± 0.009 | 0.499 ± 0.009 | 0.339 ± 0.005 | 0.491 ± 0.006 | 0.157 ± 0.016 | 0.500 ± 0.041 |
| rnaernie | 0.511 ± 0.008 | 0.526 ± 0.015 | 0.386 ± 0.009 | 0.536 ± 0.006 | 0.169 ± 0.009 | 0.494 ± 0.072 |
| rna-fm | 0.523 ± 0.009 | 0.558 ± 0.011 | 0.464 ± 0.005 | 0.599 ± 0.004 | 0.260 ± 0.012 | 0.498 ± 0.069 |
| plant-rnafm | 0.538 ± 0.008 | 0.557 ± 0.015 | 0.482 ± 0.006 | 0.619 ± 0.004 | 0.258 ± 0.012 | 0.460 ± 0.058 |
| ernierna | 0.537 ± 0.007 | 0.567 ± 0.016 | 0.508 ± 0.004 | 0.646 ± 0.005 | 0.300 ± 0.016 | 0.503 ± 0.074 |
| ernierna-ss | 0.537 ± 0.008 | 0.560 ± 0.015 | 0.491 ± 0.005 | 0.624 ± 0.006 | 0.271 ± 0.014 | 0.542 ± 0.055 |
| rinalmo-giga | 0.615 ± 0.007 | 0.560 ± 0.014 | 0.525 ± 0.006 | 0.652 ± 0.004 | 0.383 ± 0.014 | 0.459 ± 0.057 |
| rinalmo-mega | 0.610 ± 0.008 | 0.552 ± 0.012 | 0.502 ± 0.005 | 0.631 ± 0.005 | 0.358 ± 0.014 | 0.514 ± 0.051 |
| rinalmo-micro | 0.598 ± 0.007 | 0.534 ± 0.010 | 0.438 ± 0.005 | 0.581 ± 0.006 | 0.326 ± 0.011 | 0.479 ± 0.048 |
| gb-rna-1b600m | 0.563 ± 0.008 | 0.616 ± 0.015 | 0.546 ± 0.005 | 0.667 ± 0.004 | 0.324 ± 0.012 | 0.478 ± 0.089 |
| gb-rna-1b600m-cds | 0.576 ± 0.008 | 0.633 ± 0.018 | 0.559 ± 0.004 | 0.675 ± 0.005 | 0.327 ± 0.015 | 0.485 ± 0.060 |
| gb-rna-650m | 0.559 ± 0.006 | 0.613 ± 0.014 | 0.535 ± 0.005 | 0.667 ± 0.005 | 0.310 ± 0.015 | 0.499 ± 0.062 |
| gb-rna-650m-cds | 0.562 ± 0.008 | 0.632 ± 0.020 | 0.553 ± 0.006 | 0.674 ± 0.006 | 0.318 ± 0.014 | 0.496 ± 0.036 |
| helix-mrna | 0.533 ± 0.005 | 0.537 ± 0.014 | 0.428 ± 0.005 | 0.557 ± 0.007 | 0.182 ± 0.015 | 0.496 ± 0.063 |
| utrbert-3mer | 0.519 ± 0.009 | 0.591 ± 0.016 | 0.535 ± 0.004 | 0.643 ± 0.006 | 0.248 ± 0.011 | 0.514 ± 0.056 |
| utrbert-4mer | 0.520 ± 0.008 | 0.597 ± 0.013 | 0.532 ± 0.005 | 0.648 ± 0.006 | 0.249 ± 0.016 | 0.511 ± 0.059 |
| utrbert-5mer | 0.513 ± 0.007 | 0.591 ± 0.009 | 0.524 ± 0.004 | 0.637 ± 0.006 | 0.220 ± 0.019 | 0.415 ± 0.053 |
| utrbert-6mer | 0.511 ± 0.009 | 0.589 ± 0.016 | 0.521 ± 0.005 | 0.633 ± 0.006 | 0.248 ± 0.021 | 0.511 ± 0.063 |
| mrna-fm | 0.573 ± 0.008 | 0.625 ± 0.014 | 0.547 ± 0.005 | 0.693 ± 0.007 | 0.326 ± 0.008 | 0.266 ± 0.074 |
| codonbert | 0.589 ± 0.007 | 0.654 ± 0.015 | 0.549 ± 0.005 | 0.686 ± 0.007 | 0.311 ± 0.015 | 0.301 ± 0.070 |
| utrlm-mrl | 0.526 ± 0.007 | 0.524 ± 0.012 | 0.378 ± 0.004 | 0.534 ± 0.005 | 0.216 ± 0.011 | 0.386 ± 0.062 |
| utrlm-te-el | 0.529 ± 0.007 | 0.537 ± 0.014 | 0.423 ± 0.005 | 0.550 ± 0.004 | 0.211 ± 0.015 | 0.437 ± 0.063 |
| omnigenome-186m | 0.558 ± 0.008 | 0.627 ± 0.019 | 0.534 ± 0.006 | 0.669 ± 0.004 | 0.279 ± 0.010 | 0.521 ± 0.040 |
| omnigenome-52m | 0.557 ± 0.010 | 0.627 ± 0.018 | 0.535 ± 0.004 | 0.674 ± 0.006 | 0.295 ± 0.007 | 0.540 ± 0.070 |
| splicebert-h-510nt | 0.509 ± 0.007 | 0.563 ± 0.016 | 0.465 ± 0.004 | 0.595 ± 0.005 | 0.214 ± 0.011 | 0.572 ± 0.051 |
| splicebert-v-1024nt | 0.545 ± 0.006 | 0.612 ± 0.019 | 0.524 ± 0.003 | 0.656 ± 0.005 | 0.289 ± 0.010 | 0.514 ± 0.056 |
| splicebert-v-510nt | 0.537 ± 0.006 | 0.610 ± 0.017 | 0.525 ± 0.004 | 0.660 ± 0.005 | 0.263 ± 0.014 | 0.550 ± 0.047 |
| mrnabert | 0.614 ± 0.006 | 0.645 ± 0.021 | 0.586 ± 0.005 | 0.700 ± 0.004 | 0.357 ± 0.014 | 0.464 ± 0.039 |
| orthrus-base-4 | 0.612 ± 0.007 | 0.639 ± 0.014 | 0.537 ± 0.007 | 0.674 ± 0.006 | 0.347 ± 0.011 | 0.544 ± 0.061 |
| orthrus-large-4 | 0.617 ± 0.006 | 0.667 ± 0.013 | 0.563 ± 0.006 | 0.713 ± 0.005 | 0.404 ± 0.011 | 0.525 ± 0.064 |
| orthrus-large-6 | 0.623 ± 0.007 | 0.683 ± 0.016 | 0.666 ± 0.005 | 0.724 ± 0.005 | 0.407 ± 0.011 | 0.513 ± 0.089 |
| orthrus+mlm | 0.645 ± 0.006 | 0.694 ± 0.017 | 0.692 ± 0.004 | 0.751 ± 0.003 | 0.439 ± 0.010 | 0.633 ± 0.047 |

The foundation models lead on eight of 13 tasks in Table 2. The supervised CNN remains the best model on MRL-MPRA, APA, IRES and MRL-HL-Pair, and the naive baseline on miRNA-Binding. To examine the role of dataset size, we compared the standardized performance gap between the best foundation model and the CNN against log dataset size. The foundation model advantage tends to shrink as datasets grow (Pearson *r* = −0.50, *p* = 0.08; Spearman *ρ* = −0.57, *p* = 0.04) and reverses on the two largest datasets, MRL-MPRA and APA. However, this comparison is partly confounded, as the largest datasets are also the shortest and most synthetic, and MRL-HL-Pair, the smallest dataset, is an exception.

We note that Evo2 40B is not included in the main results because it performed poorly in our evaluation despite extensive troubleshooting. Evo2 20B was derived from Evo2 40B by removing later layers that showed abnormally high activation norms while contributing little to predictive loss (https://github.com/garykbrixi/evo2-20b-readme); we speculate that these layers may affect embeddings extracted from the full model. For completeness, we report its results alongside other Evo2 variants in Table 8.

### H Full Zero-shot Probing Results

We conduct zero-shot probing by extracting sequence embeddings from each model (as was done for linear-probing), computing the difference between the embeddings of the mutated and wild-type sequences, taking the L2 norm of this difference, and directly computing the AUPRC and AUROC against the pathogenic/benign labels in the TraitGym [53] and UTR variant datasets [54]. This approach was benchmarked in the original TraitGym paper [53] and was effective for a subset of the self-supervised models in attaining the best zero-shot performance. Future work could explore the use of other techniques for zero-shot variant effect prediction for mRNA such as the log-likelihood ratio, cosine distance, or inner product approaches benchmarked in [53]. We report zero-shot AUPRC for the best representative variant per model family in Figure 9, coloured by its ratio to the AUPRC of a random predictor (the fraction of pathogenic variants), and for all model variants in Table 12.

**Table 12:** Full zero-shot probing results.

| Model | **VEP TG Complex**<br>AUPRC | **VEP TG Mendelian**<br>AUPRC | **VEP UTR 5’**<br>AUPRC | **VEP UTR 3’**<br>AUPRC |
|---|---|---|---|---|
| naive-4 | 0.110 | 0.109 | 0.725 | 0.287 |
| naive-6 | 0.110 | 0.109 | 0.725 | 0.287 |
| naive-mamba | 0.113 | 0.120 | 0.833 | 0.598 |
| enformer | 0.176 | 0.198 | 0.911 | 0.684 |
| borzoi | 0.148 | 0.173 | 0.846 | 0.592 |
| flashzoi | 0.172 | 0.264 | 0.912 | 0.726 |
| alphagenome | 0.194 | 0.271 | 0.840 | 0.820 |
| evo-1-131k-base | 0.098 | 0.119 | 0.855 | 0.565 |
| evo-1-8k-base | 0.096 | 0.121 | 0.795 | 0.593 |
| evo-1.5-8k-base | 0.123 | 0.107 | 0.843 | 0.525 |
| dnabert-s | 0.123 | 0.121 | 0.778 | 0.555 |
| dnabert2 | 0.099 | 0.110 | 0.796 | 0.261 |
| hyenadna-large-1m | 0.097 | 0.119 | 0.827 | 0.591 |
| hyenadna-medium-160k | 0.116 | 0.110 | 0.800 | 0.621 |
| hyenadna-medium-450k | 0.106 | 0.115 | 0.819 | 0.599 |
| hyenadna-small-32k | 0.104 | 0.125 | 0.842 | 0.597 |
| hyenadna-tiny-16k-d128 | 0.111 | 0.119 | 0.782 | 0.558 |
| nt-2.5b-1000g | 0.106 | 0.129 | 0.861 | 0.568 |
| nt-2.5b-multi-species | 0.120 | 0.146 | 0.886 | 0.718 |
| nt-500m-1000g | 0.100 | 0.128 | 0.863 | 0.709 |
| nt-500m-human-ref | 0.112 | 0.161 | 0.879 | 0.712 |
| nt-v2-100m-multi-species | 0.134 | 0.128 | 0.861 | 0.721 |
| nt-v2-250m-multi-species | 0.111 | 0.138 | 0.826 | 0.793 |
| nt-v2-500m-multi-species | 0.112 | 0.174 | 0.864 | 0.684 |
| nt-v2-50m-multi-species | 0.114 | 0.138 | 0.874 | 0.634 |
| generanno-eukaryote-0.5b-base | 0.133 | 0.111 | 0.811 | 0.512 |
| generanno-eukaryote-1.2b-cds-annotator-preview | 0.135 | 0.143 | 0.871 | 0.841 |
| generanno-prokaryote-0.5b-base | 0.112 | 0.145 | 0.839 | 0.700 |
| generanno-prokaryote-0.5b-cds-annotator | 0.113 | 0.130 | 0.868 | 0.558 |
| nt-v3-100M-post | 0.178 | 0.223 | 0.920 | 0.837 |
| nt-v3-100M-pre | 0.121 | 0.147 | 0.868 | 0.565 |
| nt-v3-650M-post | 0.156 | 0.188 | 0.911 | 0.754 |
| nt-v3-650M-pre | 0.114 | 0.109 | 0.809 | 0.666 |
| nt-v3-8M-pre | 0.127 | 0.107 | 0.858 | 0.576 |
| generator-eukaryote-1.2b-base | 0.107 | 0.173 | 0.915 | 0.768 |
| generator-eukaryote-3b-base | 0.111 | 0.274 | 0.930 | 0.731 |
| generator-v2-eukaryote-1.2b-base | 0.120 | 0.208 | 0.922 | 0.772 |
| generator-v2-eukaryote-3b-base | 0.112 | 0.252 | 0.940 | 0.771 |
| generator-v2-prokaryote-1.2b-base | 0.103 | 0.127 | 0.871 | 0.714 |
| generator-v2-prokaryote-3b-base | 0.106 | 0.112 | 0.864 | 0.665 |
| evo2-1b-base | 0.115 | 0.133 | 0.908 | 0.792 |
| evo2-20b | 0.108 | 0.136 | 0.899 | 0.772 |
| evo2-7b | 0.113 | 0.126 | 0.897 | 0.795 |
| evo2-7b-262k | 0.110 | 0.127 | 0.908 | 0.785 |
| evo2-7b-base | 0.107 | 0.128 | 0.896 | 0.770 |
| rnabert | 0.096 | 0.160 | 0.880 | 0.618 |
| rnamsm | 0.118 | 0.124 | 0.863 | 0.647 |
| rnaernie | 0.114 | 0.131 | 0.862 | 0.664 |
| rna-fm | 0.109 | 0.122 | 0.865 | 0.685 |
| ernierna | 0.123 | 0.129 | 0.892 | 0.721 |
| ernierna-ss | 0.112 | 0.119 | 0.864 | 0.728 |
| rinalmo-giga | 0.120 | 0.127 | 0.887 | 0.693 |
| rinalmo-mega | 0.122 | 0.119 | 0.870 | 0.680 |
| rinalmo-micro | 0.117 | 0.119 | 0.845 | 0.696 |
| gb-rna-1b600m | 0.129 | 0.126 | 0.851 | 0.650 |
| gb-rna-1b600m-cds | 0.113 | 0.126 | 0.873 | 0.603 |
| gb-rna-650m | 0.129 | 0.125 | 0.864 | 0.679 |
| gb-rna-650m-cds | 0.114 | 0.133 | 0.876 | 0.678 |
| plant-rnafm | 0.123 | 0.121 | 0.830 | 0.662 |
| helix-mrna | 0.106 | 0.118 | 0.876 | 0.640 |
| utrbert-3mer | 0.128 | 0.118 | 0.880 | 0.629 |
| utrbert-4mer | 0.126 | 0.149 | 0.881 | 0.615 |
| utrbert-5mer | 0.115 | 0.136 | 0.867 | 0.624 |
| utrbert-6mer | 0.118 | 0.128 | 0.850 | 0.661 |
| mrna-fm | – | – | – | – |
| codonbert | – | – | – | – |
| utrlm-mrl | 0.115 | 0.121 | 0.846 | 0.599 |
| utrlm-te-el | 0.110 | 0.121 | 0.868 | 0.596 |
| omnigenome-186m | 0.106 | 0.114 | 0.837 | 0.634 |
| omnigenome-52m | 0.115 | 0.115 | 0.812 | 0.598 |
| splicebert-h-510nt | 0.116 | 0.121 | 0.862 | 0.669 |
| splicebert-v-1024nt | 0.123 | 0.124 | 0.876 | 0.677 |
| splicebert-v-510nt | 0.130 | 0.124 | 0.886 | 0.711 |
| mrnabert | 0.099 | 0.263 | 0.883 | 0.601 |
| orthrus-base-4 | 0.113 | 0.130 | 0.846 | 0.752 |
| orthrus-large-4 | 0.107 | 0.121 | 0.864 | 0.727 |
| orthrus-large-6 | 0.120 | 0.117 | 0.910 | 0.561 |
| orthrus+mlm | 0.115 | 0.151 | 0.906 | 0.738 |

**Figure 9:**
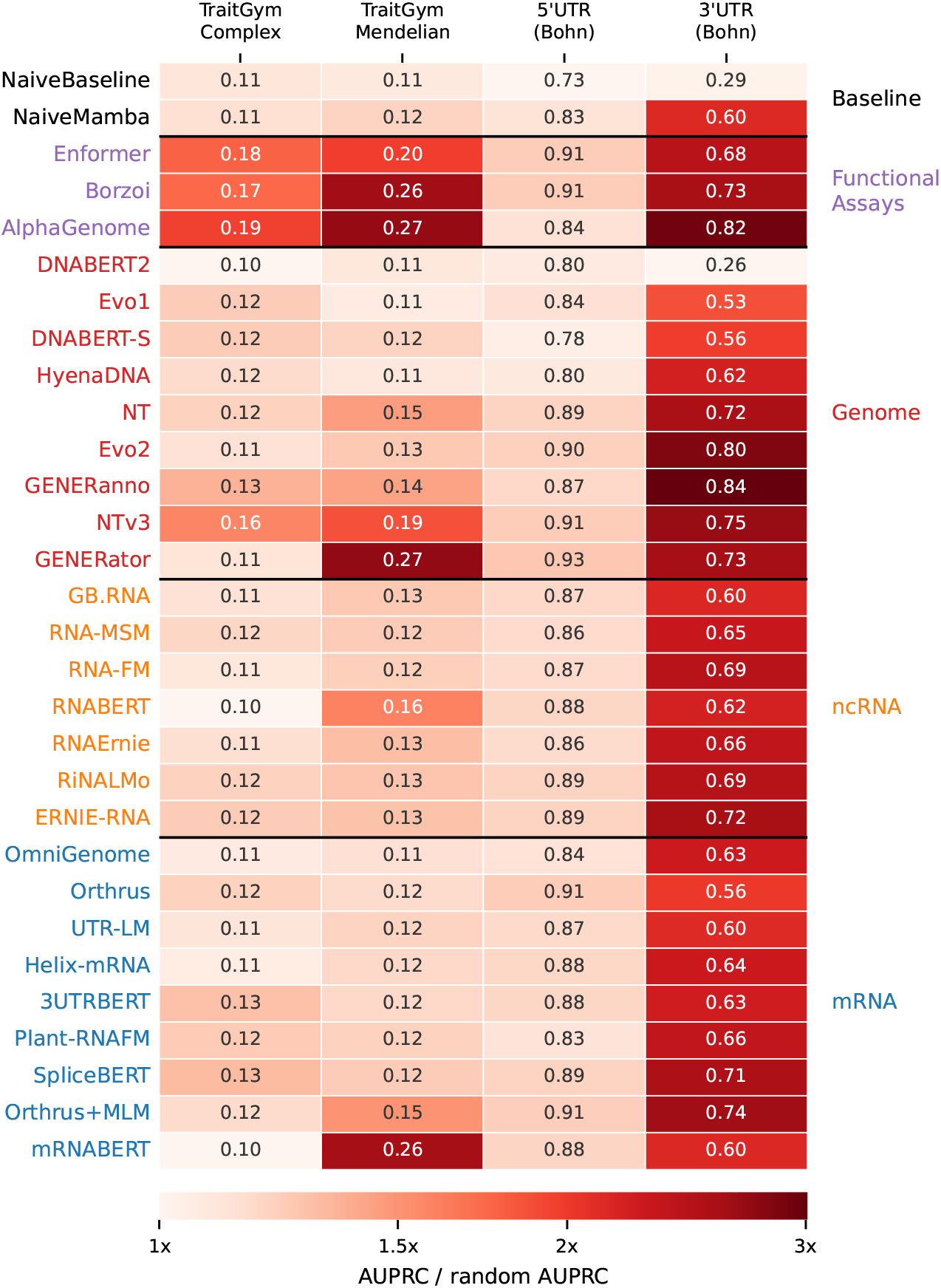
Zero-shot variant effect prediction for best performing models from Table 2. CodonBERT and mRNA-FM embed only the CDS, which none of these variants change, so they are omitted.

**Figure 10:**
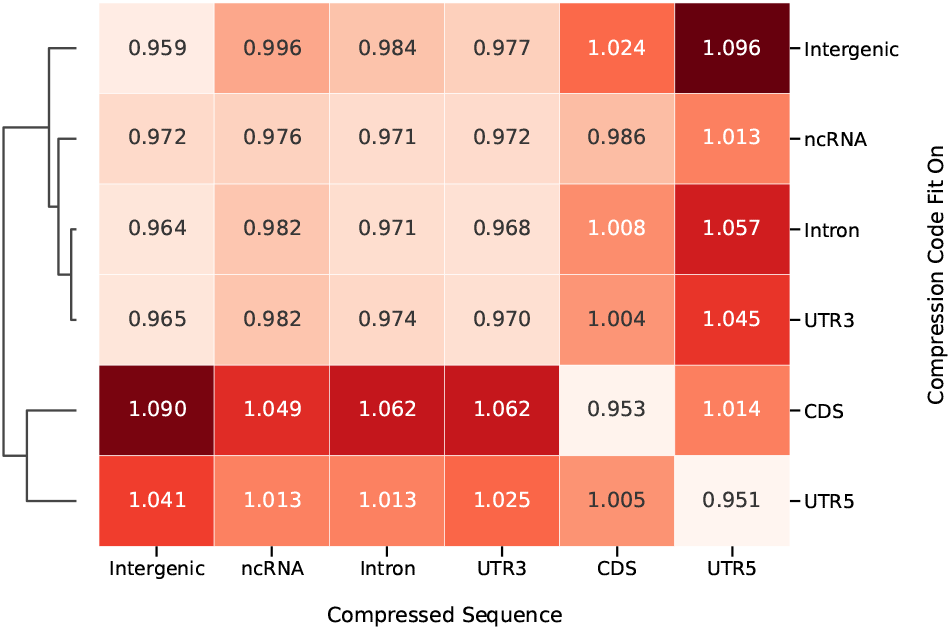
Cross-compression test scores by compression source.

**Figure 11:**
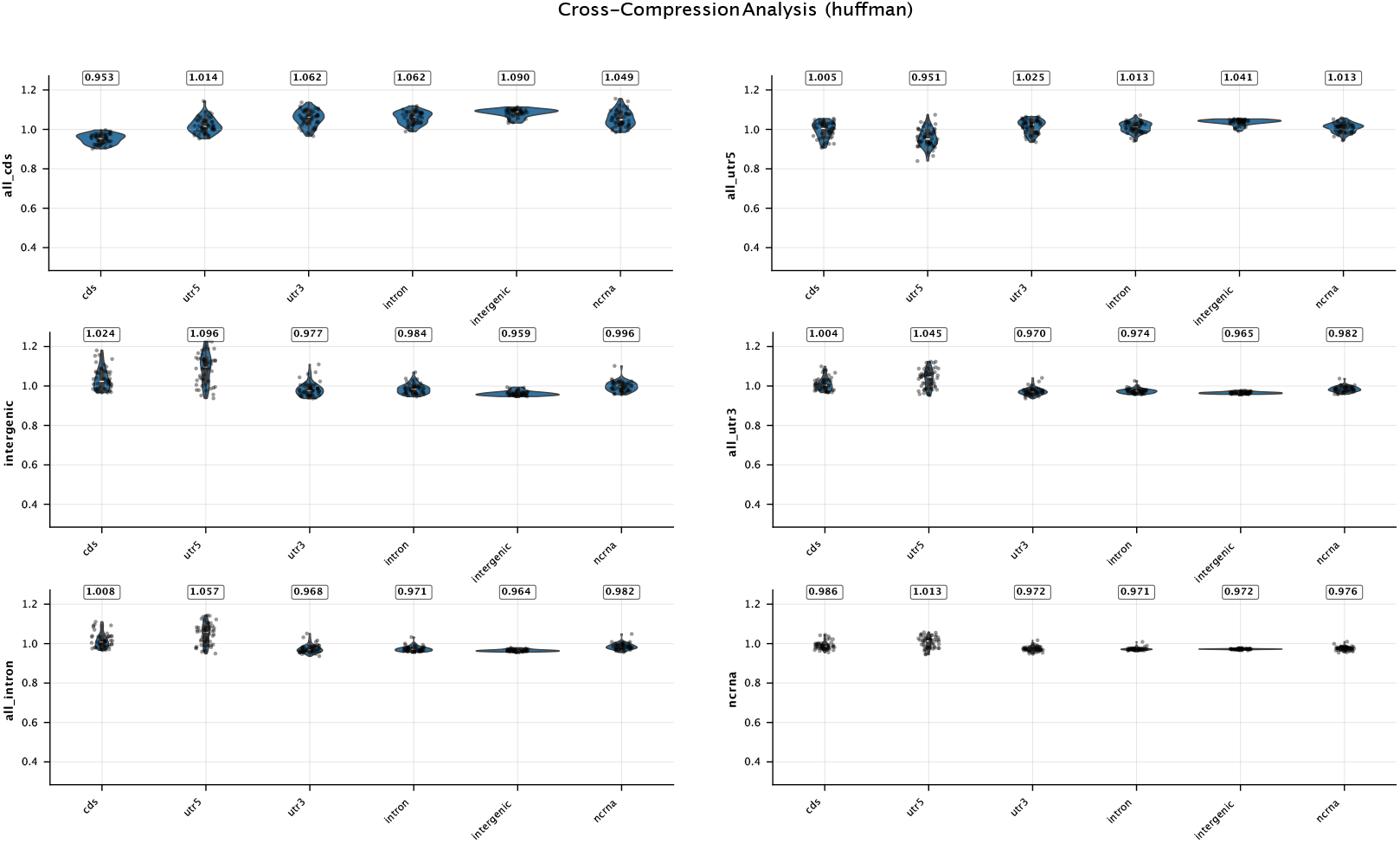
The raw distributions of 50 sequence data points compressed across Huffman codes fit to different reference sequences. 5’ UTR values demonstrate higher variability than the rest.

### I Joint Objective Experimental Setup

We perform an ablation study on the mixture of contrastive learning (CL) and masked language modelling (MLM) losses using the Orthrus model as a controlled architecture. Here, we use the architecture from the orthrus-large-6 model, which is a Mamba model with six layers and a hidden dimension of 512. See the Orthrus paper for further optimizer hyperparameters, which we matched exactly. We use a subset of the Orthrus pre-training dataset, which uses RefSeq annotations for ten species to generate positive pairs for contrastive learning. In total, this results in approximately 1M mRNA sequences.

Because the two losses differ in scale, we describe each joint model by its CL weighting 1 *− α* in the equation in Section 4.1, taken as the approximate share of the converged loss contributed by the CL term, and set the raw loss weights so that this share ranges from 0 (MLM-only) to 1 (CL-only) in steps of 0.1. For each of these 11 settings, we pre-train three models using different seeds, for a total of 33 models. Each Orthrus pre-training run took approximately 16 hours using 4 H100 GPUs. We then select the model with the highest mean validation z-score across the 11 benchmark datasets used for selection (APA and IRES are held out), computed with the same aggregation and data splitting seeds as our other overall metrics. The selected model has a CL weighting of 0.1 (*α* = 0.9). The shaded areas in Figure 3 (right) show the standard error of the mean validation z-score over the three seeds of each setting.

### J Genomic Data Compression

We define the Compression Ratio *CR* for a given sequence *s* using a specific compression model *M*. This model *M* (e.g., a Huffman codebook) is typically derived from training on a representative set of sequences from a particular source distribution (e.g., CDS, 5’ UTRs). The *CR* quantifies the efficiency of the compression by comparing the total number of bits required to represent the sequence using model *M* to its original size.

Let the sequence *s* consist of *N* = |*s*| tokens (e.g., k-mers), denoted as *s*^1^, *s*^2^, …, *s*^*N*^, *L*_*M*_ (*s*^*k*^) be the code length (in bits) assigned to the kth token *s*^*k*^ by the model *M*. Let *b*_*original*_ be the number of bits used to represent each token in the original, uncompressed fixed-length format (e.g., 2×kmer length for DNA k-mers). The Compression Ratio is then:

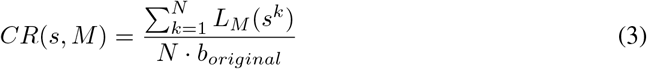

The numerator represents the total compressed size of the sequence *s* in bits when encoded using model *M*. The denominator represents the total original size of the sequence in bits. A lower *CR* indicates better compression, reflecting higher statistical regularity in the sequence *s* as captured by the model *M*. This normalized metric facilitates comparisons of compressibility across different sequences and data domains.

When comparing two different compression models, *M*_*l*_ and *M*_*m*_, fit on distinct data sources (e.g., source *l* and source *m*), we can assess their relative effectiveness on a specific sequence *s*. If *s* originates from the same source distribution as the training data for *M*_*m*_, the difference:

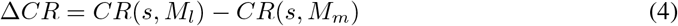

quantifies the loss in compression efficacy when using a model trained on a different data source *M*_*l*_ compared to the model trained on the native source *M*_*m*_. A positive Δ*CR* indicates that model *M*_*l*_ is less efficient for sequence *s*, suggesting a quantifiable difference between the underlying statistical distributions of the data sources *l* and *m*.

#### Algorithm 1

Huffman Coding Pseudocode

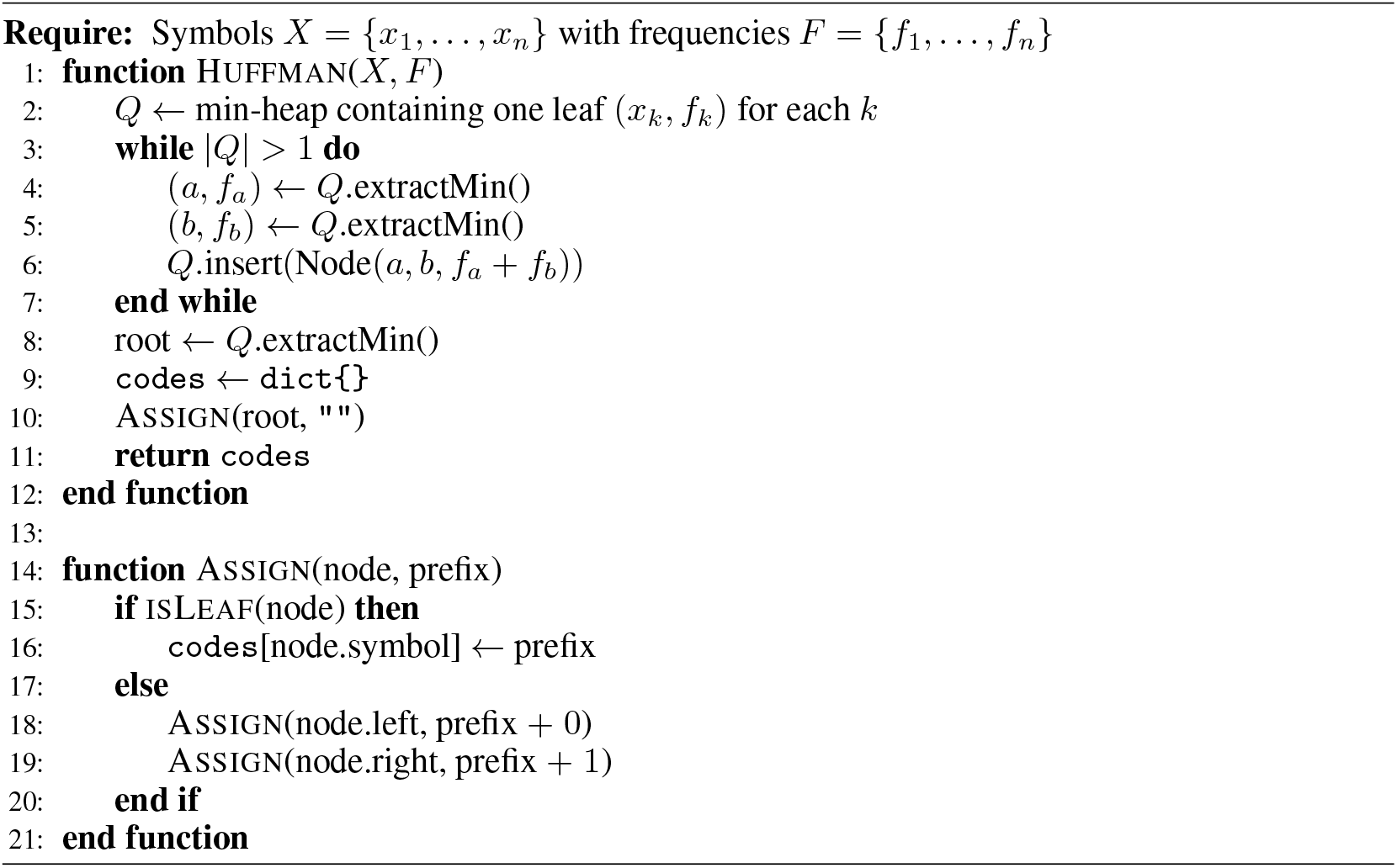

For the compression algorithm we utilize Huffman coding, shown in Algorithm 1, which is a widely used algorithm for lossless data compression. Its strategy is to assign shorter binary codes to more frequent symbols and longer codes to less frequent ones, thereby minimizing the average code length for a given text or data sample.

Specifically, Huffman compression relies on empirically estimating the probability of discrete events from a given data sample. Based on these estimated probabilities, the algorithm assigns variable length binary codes to each event. The length of these codes is inversely proportional to the symbol’s probability: more frequent symbols receive shorter codes. For an event *x*_*i*_ with probability *p*(*x*_*i*_), the optimal code length is *−*log_2_ *p*(*x*_*i*_) bits. Huffman coding aims to construct a prefix code that approaches this theoretical optimum in terms of average code length.

Let *P*_*M*_ be the probability distribution of tokens (symbols) implicitly learned by a compression model *M* during its training (e.g., from the frequencies of k-mers in a training sequence). The average code length assigned by model *M* to tokens from this distribution *P*_*M*_, denoted *L*_*M*_, satisfies *H*(*P*_*M*_) *≤ L*_*M*_ *< H*(*P*_*M*_) + 1, where *H*(*P*_*M*_) is the Shannon entropy of the source distribution *P*_*M*_ :

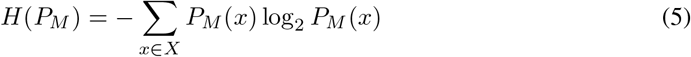

Here, *X* is the set of all possible tokens. This indicates that Huffman coding is nearly optimal for the data distribution it was trained on.

Similarly to how the compression ratio *CR* of a sequence *s* has a close connection to the underlying entropy of a distribution, we can draw a similar connection to measuring cross entropy between two distributions. This is done by applying a compression model *M* fit on *P*_*M*_ to compress a sequence *s*_*T*_ sampled from an underlying probability distribution *P*_*T*_.

Consider two probability distributions defined over the same sample space *P*_*M*_ and *P*_*T*_. Assume sequence *s*_*T*_ is drawn from *P*_*T*_. When we use a compression model M (which was fit on *P*_*M*_) to compress *s*_*T*_, the average number of bits per token used to encode *s*_*T*_ is

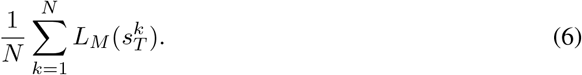

This empirical average code length is an estimate of the cross-entropy between the true distribution *P*_*T*_ and the model’s distribution *P*_*M*_ :

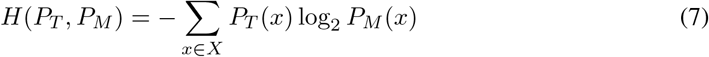

The term *L*_*M*_ (*s*^*k*^) represents the code length assigned by model *M* to the specific token *s*^*k*^. In the case of using Huffman coding compression *P*_*M*_ (*x*) are the probabilities used to construct the Huffman codes in model *M*, then *L*_*M*_ (*x*) ≈ − log_2_ *P*_*M*_ (*x*) for each token *x*.

The numerator of the Compression Ratio *CR* (Equation 3) is the total compressed size of sequence s using model M. Thus, 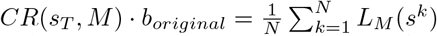, which is the empirical average bits per token. Therefore, the Compression Ratio *CR*(*s*_*T*_, *M*) is directly proportional to this empirical estimate of *H*(*P*_*T*_, *P*_*M*_).

It is important to note that this analysis, based on token frequencies and Huffman coding, has inherent limitations. The choice of token size (e.g., k-mer length) is critical: larger tokens can capture more local context but may result in an exponentially larger sample space. This can make it challenging to empirically estimate the true probability distributions from finite training data, potentially leading to inaccurate frequency counts for rare events. In addition, certain structures within the genome have a specific repetition order such as an open reading frame in coding sequences consisting of a sequence of codons composed of three nucleotides each. We use k-mer length of six to capture this regularity.

Furthermore, by focusing on individual token frequencies, this approach largely ignores global sequence structure and long-range dependencies. Such larger-scale structures, including regulatory motifs, codon usage patterns beyond local k-mers, and repetitive elements, are known to be significant features in both coding and intergenic regions and are not directly captured by this type of local, frequency-based compression analysis.

Training data for the compression analysis was generated using GenomeKit [34] and GENCODE Basic version 47 annotations [97]. From these annotations, we isolated 3’ UTRs, 5’ UTRs, introns, and coding sequences (CDS) of protein-coding genes.

Because intergenic regions are vast, we restricted our intergenic sequence set to those on chromosome 1, which is about 8% of the human genome. This selection helped balance the corresponding length of different genomic region types in our training data, reducing potential biases in the compression model fitting that could arise (Table 13). For genes with multiple transcripts, we selected the canonical isoform using the APPRIS database [39] to ensure consistency and reduce redundancy from alternative splice variants.

**Table 13:** Distribution and characteristics of sequences with and without repeats across various genomic regions. Counts represent the number of distinct sequences. Lengths are reported in base pairs (bp). The number of bases used for estimating the compression is omitting repeats.

| Genomic Region | Num. Sequences with Repeats | Total Length with Repeats (bp) | Num. Sequences without Repeats | Total Length without Repeats (bp) |
|---|---|---|---|---|
| CDS | 3,706 | 4,927,801 | 3,121 | 3,883,277 |
| Intron | 2,922 | 91,533,879 | 415 | 569,014 |
| UTR3 | 3,506 | 5,271,453 | 2,241 | 1,902,431 |
| UTR5 | 3,499 | 629,402 | 3,127 | 496,448 |
| Intergenic(chr1) | 196 | 13,640,727 | 26,166 | 15,732,853 |
| ncRNA | 38,016 | 33,615,313 | 14,648 | 9,549,965 |

When preparing training data for each category, we excluded a hold-out set of 50 unique sequences of variable lengths. These sequences were never seen by the compression algorithm during its fitting stage and were used as a test set. For the Huffman compression algorithm, all sequences (training and test) were first divided into non-overlapping k-mers of length six. Sequences were then truncated at their 3’ end to ensure their total length was a multiple of six, losing at most five nucleotides per sequence. This preprocessing was crucial to keep the reading frame intact when sequences within each category were combined for frequency analysis by the compression algorithm.

Finally, to check if repetitive elements affected our findings, we conducted a parallel analysis where repeat regions, identified by RepeatMasker [98], were excluded from all genomic categories. The results from this repeat-masked analysis were very similar to those in the main paper, suggesting common repeats did not significantly affect our conclusions (Figure 12).

**Figure 12:**
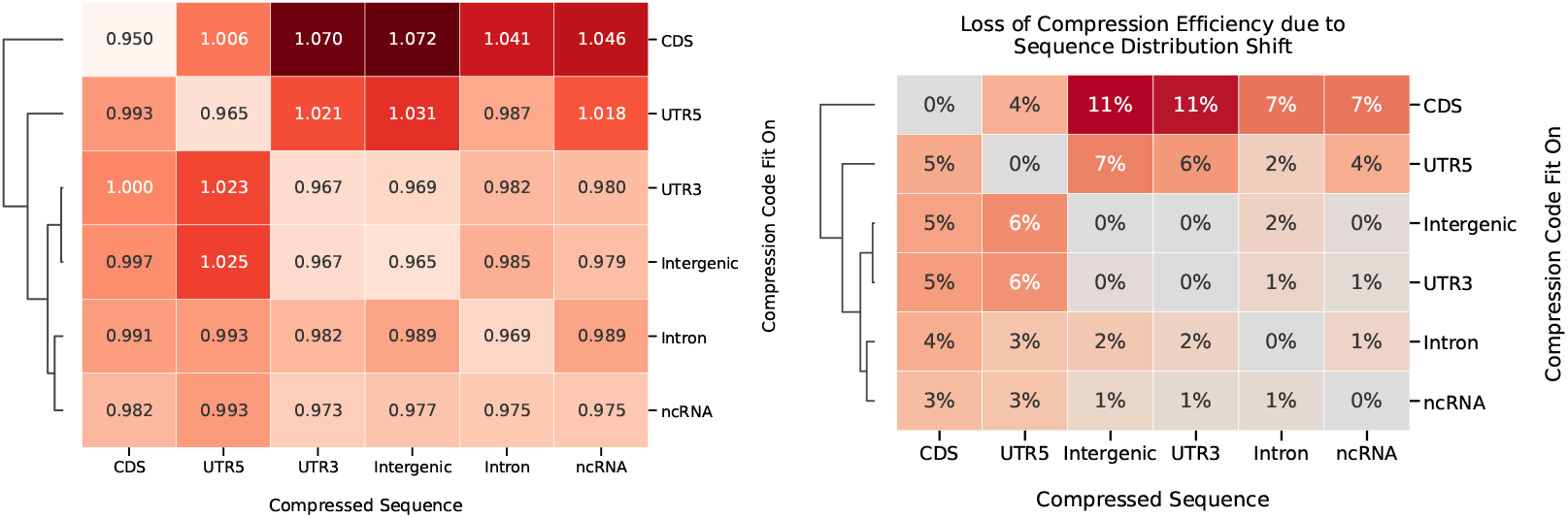
**Left:** Compression ratios indicating the percentage of size reduction associated with applying Huffman coding algorithm on the source data with k-mer length set to 6. All regions that contain repeats as indicated by repeat masker are omitted. **Right:** Cross-compression test scores by compression source. Numbers show percent increase in compression ratio while omitting genomic regions containing repeats.

#### NeurIPS Paper Checklist

##### 1. Claims

Question: Do the main claims made in the abstract and introduction accurately reflect the paper’s contributions and scope?

Answer: [Yes]

Justification: Main claims include a benchmarking suite on mRNA datasets and prediction tasks using nucleotide foundation models, which are included as code and evaluation results. The abstract claims an assessment of parameter scaling, compression, and data splitting, which are reported in sections 5, 5.2, and 5.3 respectively. Finally, the abstract claims a new model trained using mRNABench insights, which is reported in section 5.1.

Guidelines:

- The answer [N/A] means that the abstract and introduction do not include the claims made in the paper.
- The abstract and/or introduction should clearly state the claims made, including the contributions made in the paper and important assumptions and limitations. A [No] or [N/A] answer to this question will not be perceived well by the reviewers.
- The claims made should match theoretical and experimental results, and reflect how much the results can be expected to generalize to other settings.
- It is fine to include aspirational goals as motivation as long as it is clear that these goals are not attained by the paper.

##### 2. Limitations

Question: Does the paper discuss the limitations of the work performed by the authors? Answer: [Yes]

Justification: The paper contains a limitations section, highlighting issues such as models which we could not get to converge, models which were omitted from evaluation, and tasks which were not included.

Guidelines:

- The answer [N/A] means that the paper has no limitation while the answer [No] means that the paper has limitations, but those are not discussed in the paper.
- The authors are encouraged to create a separate “Limitations” section in their paper.
- The paper should point out any strong assumptions and how robust the results are to violations of these assumptions (e.g., independence assumptions, noiseless settings, model well-specification, asymptotic approximations only holding locally). The authors should reflect on how these assumptions might be violated in practice and what the implications would be.
- The authors should reflect on the scope of the claims made, e.g., if the approach was only tested on a few datasets or with a few runs. In general, empirical results often depend on implicit assumptions, which should be articulated.
- The authors should reflect on the factors that influence the performance of the approach. For example, a facial recognition algorithm may perform poorly when image resolution is low or images are taken in low lighting. Or a speech-to-text system might not be used reliably to provide closed captions for online lectures because it fails to handle technical jargon.
- The authors should discuss the computational efficiency of the proposed algorithms and how they scale with dataset size.
- If applicable, the authors should discuss possible limitations of their approach to address problems of privacy and fairness.
- While the authors might fear that complete honesty about limitations might be used by reviewers as grounds for rejection, a worse outcome might be that reviewers discover limitations that aren’t acknowledged in the paper. The authors should use their best judgment and recognize that individual actions in favor of transparency play an important role in developing norms that preserve the integrity of the community. Reviewers will be specifically instructed to not penalize honesty concerning limitations.

##### 3. Theory assumptions and proofs

Question: For each theoretical result, does the paper provide the full set of assumptions and a complete (and correct) proof?

Answer: [N/A]

Justification: Paper does not include theoretical results. Guidelines:

- The answer [N/A] means that the paper does not include theoretical results.
- All the theorems, formulas, and proofs in the paper should be numbered and cross-referenced.
- All assumptions should be clearly stated or referenced in the statement of any theorems.
- The proofs can either appear in the main paper or the supplemental material, but if they appear in the supplemental material, the authors are encouraged to provide a short proof sketch to provide intuition.
- Inversely, any informal proof provided in the core of the paper should be complemented by formal proofs provided in appendix or supplemental material.
- Theorems and Lemmas that the proof relies upon should be properly referenced.

##### 4. Experimental result reproducibility

Question: Does the paper fully disclose all the information needed to reproduce the main experimental results of the paper to the extent that it affects the main claims and/or conclusions of the paper (regardless of whether the code and data are provided or not)?

Answer: [Yes]

Justification: The submission is accompanied by code that allows the main benchmark evaluations to be reproduced. All datasets and benchmarked models are publicly accessible. Furthermore, the experimental setups are reported in Appendix C, and data processing pipelines reported in Appendix B. Supervised baseline training hyperparameters were reported in Appendix A. All results are computed using fixed seeds, allowing exact reproducibility.

Guidelines:

- The answer [N/A] means that the paper does not include experiments.
- If the paper includes experiments, a [No] answer to this question will not be perceived well by the reviewers: Making the paper reproducible is important, regardless of whether the code and data are provided or not.
- If the contribution is a dataset and/or model, the authors should describe the steps taken to make their results reproducible or verifiable.
- Depending on the contribution, reproducibility can be accomplished in various ways. For example, if the contribution is a novel architecture, describing the architecture fully might suffice, or if the contribution is a specific model and empirical evaluation, it may be necessary to either make it possible for others to replicate the model with the same dataset, or provide access to the model. In general. releasing code and data is often one good way to accomplish this, but reproducibility can also be provided via detailed instructions for how to replicate the results, access to a hosted model (e.g., in the case of a large language model), releasing of a model checkpoint, or other means that are appropriate to the research performed.
- While NeurIPS does not require releasing code, the conference does require all submissions to provide some reasonable avenue for reproducibility, which may depend on the nature of the contribution. For example

a. If the contribution is primarily a new algorithm, the paper should make it clear how to reproduce that algorithm.
b. If the contribution is primarily a new model architecture, the paper should describe the architecture clearly and fully.
c. If the contribution is a new model (e.g., a large language model), then there should either be a way to access this model for reproducing the results or a way to reproduce the model (e.g., with an open-source dataset or instructions for how to construct the dataset).
d. We recognize that reproducibility may be tricky in some cases, in which case authors are welcome to describe the particular way they provide for reproducibility. In the case of closed-source models, it may be that access to the model is limited in some way (e.g., to registered users), but it should be possible for other researchers to have some path to reproducing or verifying the results.

##### 5. Open access to data and code

Question: Does the paper provide open access to the data and code, with sufficient instructions to faithfully reproduce the main experimental results, as described in supplemental material?

Answer: [Yes]

Justification: The code and datasets that are included as part of the benchmark are hosted with open access. A detailed README describes how users can install and run benchmarking results from end-to-end. The paper’s appendices also describe how datasets are processed, and raw dataset sources.

Guidelines:

- The answer [N/A] means that paper does not include experiments requiring code.
- Please see the NeurIPS code and data submission guidelines (https://neurips.cc/public/guides/CodeSubmissionPolicy) for more details.
- While we encourage the release of code and data, we understand that this might not be possible, so [No] is an acceptable answer. Papers cannot be rejected simply for not including code, unless this is central to the contribution (e.g., for a new open-source benchmark).
- The instructions should contain the exact command and environment needed to run to reproduce the results. See the NeurIPS code and data submission guidelines (https://neurips.cc/public/guides/CodeSubmissionPolicy) for more details.
- The authors should provide instructions on data access and preparation, including how to access the raw data, preprocessed data, intermediate data, and generated data, etc.
- The authors should provide scripts to reproduce all experimental results for the new proposed method and baselines. If only a subset of experiments are reproducible, they should state which ones are omitted from the script and why.
- At submission time, to preserve anonymity, the authors should release anonymized versions (if applicable).
- Providing as much information as possible in supplemental material (appended to the paper) is recommended, but including URLs to data and code is permitted.

##### 6. Experimental setting/details

Question: Does the paper specify all the training and test details (e.g., data splits, hyperparameters, how they were chosen, type of optimizer) necessary to understand the results?

Answer: [Yes]

Justification: Appendices A-C describe the various data splitting, hyperparameter selection, and optimization strategies used in the evaluation.

Guidelines:

- The answer [N/A] means that the paper does not include experiments.
- The experimental setting should be presented in the core of the paper to a level of detail that is necessary to appreciate the results and make sense of them.
- The full details can be provided either with the code, in appendix, or as supplemental material.

##### 7. Experiment statistical significance

Question: Does the paper report error bars suitably and correctly defined or other appropriate information about the statistical significance of the experiments?

Answer: [Yes]

Justification: The main result table 2 reports the statistical test used to determine significance, source of variability (random seeds affect split membership), and number of splits used. For visualization clarity, the 95% confidence intervals are reported in Appendix Tables 9, 10, and 11 instead of main paper. In general, where error bars are shown, source of variation is described, and statistical tests, if applicable, are described.

Guidelines:

- The answer [N/A] means that the paper does not include experiments.
- The authors should answer [Yes] if the results are accompanied by error bars, confidence intervals, or statistical significance tests, at least for the experiments that support the main claims of the paper.
- The factors of variability that the error bars are capturing should be clearly stated (for example, train/test split, initialization, random drawing of some parameter, or overall run with given experimental conditions).
- The method for calculating the error bars should be explained (closed form formula, call to a library function, bootstrap, etc.)
- The assumptions made should be given (e.g., Normally distributed errors).
- It should be clear whether the error bar is the standard deviation or the standard error of the mean.
- It is OK to report 1-sigma error bars, but one should state it. The authors should preferably report a 2-sigma error bar than state that they have a 96% CI, if the hypothesis of Normality of errors is not verified.
- For asymmetric distributions, the authors should be careful not to show in tables or figures symmetric error bars that would yield results that are out of range (e.g., negative error rates).
- If error bars are reported in tables or plots, the authors should explain in the text how they were calculated and reference the corresponding figures or tables in the text.

##### 8. Experiments compute resources

Question: For each experiment, does the paper provide sufficient information on the computer resources (type of compute workers, memory, time of execution) needed to reproduce the experiments?

Answer: [Yes]

Justification: The exact GPUs and runtimes of experiments are reported in the Appendix. Due to the relatively low methodological novelty of the paper, relatively few additional experiments in excess of reported computation was required.

Guidelines:

- The answer [N/A] means that the paper does not include experiments.
- The paper should indicate the type of compute workers CPU or GPU, internal cluster, or cloud provider, including relevant memory and storage.
- The paper should provide the amount of compute required for each of the individual experimental runs as well as estimate the total compute.
- The paper should disclose whether the full research project required more compute than the experiments reported in the paper (e.g., preliminary or failed experiments that didn’t make it into the paper).

##### 9. Code of ethics

Question: Does the research conducted in the paper conform, in every respect, with the NeurIPS Code of Ethics https://neurips.cc/public/EthicsGuidelines?

Answer: [Yes]

Justification: We believe the work in this submission conforms to the code of ethics, and does not pose any ethical risks.

Guidelines:

- The answer [N/A] means that the authors have not reviewed the NeurIPS Code of Ethics.
- If the authors answer [No], they should explain the special circumstances that require a deviation from the Code of Ethics.
- The authors should make sure to preserve anonymity (e.g., if there is a special consideration due to laws or regulations in their jurisdiction).

##### 10. Broader impacts

Question: Does the paper discuss both potential positive societal impacts and negative societal impacts of the work performed?

Answer: [N/A]

Justification: We do not feel that there are significant societal impacts that could result from this benchmark. All data and models comes from previously available sources, and its integration in this benchmark does not yield insights that we believe could be harmful.

Guidelines:

- The answer [N/A] means that there is no societal impact of the work performed.
- If the authors answer [N/A] or [No], they should explain why their work has no societal impact or why the paper does not address societal impact.
- Examples of negative societal impacts include potential malicious or unintended uses (e.g., disinformation, generating fake profiles, surveillance), fairness considerations (e.g., deployment of technologies that could make decisions that unfairly impact specific groups), privacy considerations, and security considerations.
- The conference expects that many papers will be foundational research and not tied to particular applications, let alone deployments. However, if there is a direct path to any negative applications, the authors should point it out. For example, it is legitimate to point out that an improvement in the quality of generative models could be used to generate Deepfakes for disinformation. On the other hand, it is not needed to point out that a generic algorithm for optimizing neural networks could enable people to train models that generate Deepfakes faster.
- The authors should consider possible harms that could arise when the technology is being used as intended and functioning correctly, harms that could arise when the technology is being used as intended but gives incorrect results, and harms following from (intentional or unintentional) misuse of the technology.
- If there are negative societal impacts, the authors could also discuss possible mitigation strategies (e.g., gated release of models, providing defenses in addition to attacks, mechanisms for monitoring misuse, mechanisms to monitor how a system learns from feedback over time, improving the efficiency and accessibility of ML).

##### 11. Safeguards

Question: Does the paper describe safeguards that have been put in place for responsible release of data or models that have a high risk for misuse (e.g., pre-trained language models, image generators, or scraped datasets)?

Answer: [N/A]

Justification: We do not feel that there are risks caused by this benchmark that require safeguards.

Guidelines:

- The answer [N/A] means that the paper poses no such risks.
- Released models that have a high risk for misuse or dual-use should be released with necessary safeguards to allow for controlled use of the model, for example by requiring that users adhere to usage guidelines or restrictions to access the model or implementing safety filters.
- Datasets that have been scraped from the Internet could pose safety risks. The authors should describe how they avoided releasing unsafe images.
- We recognize that providing effective safeguards is challenging, and many papers do not require this, but we encourage authors to take this into account and make a best faith effort.

##### 12. Licenses for existing assets

Question: Are the creators or original owners of assets (e.g., code, data, models), used in the paper, properly credited and are the license and terms of use explicitly mentioned and properly respected?

Answer: [Yes]

Justification: We have cited each original dataset and model. On each curated data source, we have provided a link and citation to the original paper. Similarly for each rehosted model, we have provided original attribution. The license of each dataset has been described.

Guidelines:

- The answer [N/A] means that the paper does not use existing assets.
- The authors should cite the original paper that produced the code package or dataset.
- The authors should state which version of the asset is used and, if possible, include a URL.
- The name of the license (e.g., CC-BY 4.0) should be included for each asset.
- For scraped data from a particular source (e.g., website), the copyright and terms of service of that source should be provided.
- If assets are released, the license, copyright information, and terms of use in the package should be provided. For popular datasets, paperswithcode.com/datasets has curated licenses for some datasets. Their licensing guide can help determine the license of a dataset.
- For existing datasets that are re-packaged, both the original license and the license of the derived asset (if it has changed) should be provided.
- If this information is not available online, the authors are encouraged to reach out to the asset’s creators.

##### 13. New assets

Question: Are new assets introduced in the paper well documented and is the documentation provided alongside the assets?

Answer: [Yes]

Justification: This paper introduces code for a new benchmarking framework, which is documented through in a detailed README and within the appendices of the paper submission. Assets which were used were typically listed under creative commons or similar license which did not require prior consent for integration.

Guidelines:

- The answer [N/A] means that the paper does not release new assets.
- Researchers should communicate the details of the dataset/code/model as part of their submissions via structured templates. This includes details about training, license, limitations, etc.
- The paper should discuss whether and how consent was obtained from people whose asset is used.
- At submission time, remember to anonymize your assets (if applicable). You can either create an anonymized URL or include an anonymized zip file.

##### 14. Crowdsourcing and research with human subjects

Question: For crowdsourcing experiments and research with human subjects, does the paper include the full text of instructions given to participants and screenshots, if applicable, as well as details about compensation (if any)?

Answer: [N/A]

Justification: Paper does not involve crowdsourcing or human subject research. Guidelines:

- The answer [N/A] means that the paper does not involve crowdsourcing nor research with human subjects.
- Including this information in the supplemental material is fine, but if the main contribution of the paper involves human subjects, then as much detail as possible should be included in the main paper.
- According to the NeurIPS Code of Ethics, workers involved in data collection, curation, or other labor should be paid at least the minimum wage in the country of the data collector.

##### 15. Institutional review board (IRB) approvals or equivalent for research with human subjects

Question: Does the paper describe potential risks incurred by study participants, whether such risks were disclosed to the subjects, and whether Institutional Review Board (IRB) approvals (or an equivalent approval/review based on the requirements of your country or institution) were obtained?

Answer: [N/A]

Justification: Paper does not use human subjects, and does not require IRB approval. Guidelines:

- The answer [N/A] means that the paper does not involve crowdsourcing nor research with human subjects.
- Depending on the country in which research is conducted, IRB approval (or equivalent) may be required for any human subjects research. If you obtained IRB approval, you should clearly state this in the paper.
- We recognize that the procedures for this may vary significantly between institutions and locations, and we expect authors to adhere to the NeurIPS Code of Ethics and the guidelines for their institution.
- For initial submissions, do not include any information that would break anonymity (if applicable), such as the institution conducting the review.

##### 16. Declaration of LLM usage

Question: Does the paper describe the usage of LLMs if it is an important, original, or non-standard component of the core methods in this research? Note that if the LLM is used only for writing, editing, or formatting purposes and does *not* impact the core methodology, scientific rigor, or originality of the research, declaration is not required.

Answer: [N/A]

Justification: Core methodology does not rely on LLMs. Guidelines:

- The answer [N/A] means that the core method development in this research does not involve LLMs as any important, original, or non-standard components.
- Please refer to our LLM policy in the NeurIPS handbook for what should or should not be described.

## Notes

### Summary of Updates

The manuscript has been updated with additional datasets, evaluated models, and improved analysis.

